# Joint analysis of multiply perturbed cells improves statistical power and cost efficiency in Perturb-seq

**DOI:** 10.64898/2026.07.10.737863

**Authors:** Jake Yeung, Jenille Tan, Liang Wang, Diana Wu, Sandra Melo Carlos, Jorge Kageyama, Jack Kamm, Benjamin B. Chu, Oleg Mayba, William F. Forrest, Shiqi Xie

## Abstract

Perturb-seq measures transcriptomic responses to genetic perturbations at scale, but conventional designs that enrich for one guide RNA per cell remain resource-intensive. Standard analyses discard cells carrying multiple guides, further limiting the usable yield from each experiment. Here, we characterize how incorporating these guide multiplets affects signal recovery, information loss, and cost reduction. At the highest guide burden, cells showed increased stress and suppressed cell-cycle progression. We develop PerturbMatch, a scalable statistical framework to analyze guide multiplets. Among different classes of guide multiplets, doublets and triplets recovered perturbation responses more accurately than higher-order multiplets. Across three 5000-gene Perturb-seq screens with increasing guide loading, per-cell costs decreased by up to 81% while information loss remained within 1.5-fold of the loss observed between technical replicates. In existing genome-wide Perturb-seq data, incorporating previously discarded guide multiplets increased usable cell numbers and improved statistical power. Compared with a singlet holdout set, adding guide multiplets moved signal recovery closer to the theoretical expected reproducibility. Overall, we recommend a design that intentionally includes single-guide cells, guide doublets, and guide triplets to improve cost efficiency while preserving signal recovery.

## Introduction

Pooled single-cell CRISPR-seq screens [1] (including Perturb-seq [2, 3], CRISP-seq [4], CROP-seq [5], and Mosaic-seq [6]; hereafter referred to as Perturb-seq) have become a foundational technology to measure transcriptomic responses to genetic perturbations. Recent studies show technical feasibility to scale genetic perturbations to more than 10 000 genes per condition [7, 8]. These successes have inspired efforts to map large-scale perturbation responses across cultured cell lines [8], primary cells [9], complex *in vitro* models [10], and *in vivo* tissues [11]. However, experimental designs and their accompanying analyses have not yet addressed the cost and complexity of Perturb-seq screens at these new scales. Conventional Perturb-seq designs aim to recover on average one guide RNA per cell, and discard any cell carrying multiple guides during analysis.

Previous studies show that overloading cells with single-guide RNAs (hereafter, guides) and then analyzing the transcriptomic readouts of guide multiplets can reduce costs in enhancer screens [12, 13]. In a recent 598-gene compressed Perturb-seq screen [14], comparisons between different compression strategies have favored pooling guides in cells to make guide multiplets (by increasing multiplicity of infection, MOI) over pooling cells in droplets (by increasing cell concentration). Despite these results, high-MOI Perturb-seq has not yet been systematically benchmarked as a scalable strategy for perturbation data. Key uncertainties remain. First, what systematic transcriptional artifacts are provoked by extreme guide burden? Second, can simple computational frameworks scale guide multiplet analysis across thousands of perturbations? Third, what is the quantitative trade-off between cost reduction and information loss as we increase the fraction of guide multiplets?

Here, we generate targeted and unbiased Perturb-seq screens at different levels of MOI, with a total of two million analyzed cells across nine screens. We develop PerturbMatch, a scalable statistical framework to analyze guide multiplets across thousands of perturbations and millions of cells. Using PerturbMatch, we quantify how different proportions of guide multiplets affect signal recovery. Finally, we establish a theoretical framework relating transcriptome-wide response magnitude to expected reproducibility across MOI levels.

We find that increasing guide burden eventually induces cellular stress and arrests cell cycle progression. Guide doublets and triplets recapitulate perturbation responses better than higher-order multiplets. In an untargeted Perturb-seq screen knocking down 5000 genes, we show that the high-MOI strategy can reduce costs by 81% while maintaining relative information loss within 1.5-fold of a low-MOI baseline. Finally, we perform joint analysis on the 9000-gene Perturb-seq dataset from Replogle et al. [7]. By incorporating previously discarded guide multiplets, we improve statistical power and drive signal recovery towards the theoretical expected reproducibility.

## Results

### Extreme guide burden induces stress and suppresses cell-cycle progression

To test the limits of increasing multiplicity of infection (MOI) on the guide burden across cells, we performed targeted CRISPRi Perturb-seq in the DLD-1 cell line across 45 promoters (targeting 35 genes) at six levels of MOI (normal infection at low, medium, and high MOI; spinfection at low, medium, high MOI; Methods 1.4, Figure 1A). The genes chosen here all have known function in DLD-1 and we therefore expected most to have detectable transcriptomic phenotypes. In this targeted Perturb-seq, we recovered 93 305 cells after quality control (88 287 cells with *>* 0 guides, Methods 1.6), with a median of 7700 cells per guide per 100 000 cells (normalized by 100 000 to simplify comparisons between experiments with different numbers of cells, Figure 1B, C). In all analyses, we focused only on droplets containing one cell and discarded droplets containing multiple cells. We therefore refer to single cells carrying multiple guides as “multiply perturbed cells,” “guide multiplets,” or simply “multiplets”. We assigned guides to each cell using a recent contingency table approach, called fishash [15]. At the two highest levels of MOI (spinfection medium and high), we saturated at a mean of 7.9 to 8.3 guides per cell (Figure 1D), giving a theoretical cost reduction of up to 88% (assuming a mean of 8.3 guides per cell would boost the effective number of cells by a factor of 8.3 with the same experimental budget). This limit was lower than previous high-MOI studies in K562 cells, which achieved a median of 12–25 guides [12, 13], suggesting that the intrinsic limit of guide burden varies across cell lines.

**Fig. 1.**
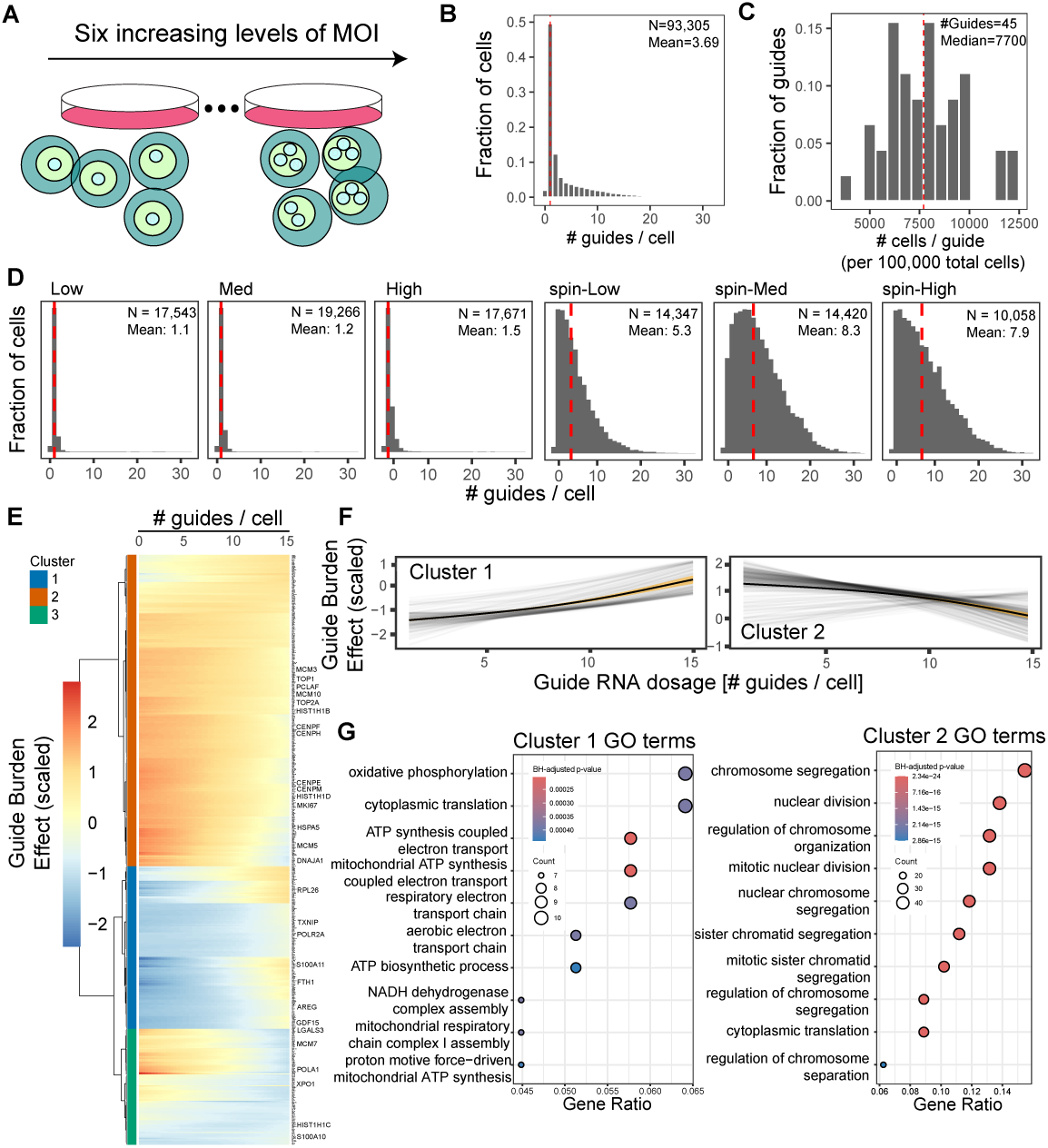
Extreme guide burden induces stress and suppresses cell-cycle progression. **A**, Experimental design for six Perturb-seq experiments with increasing multiplicity of infection (MOI). Cartoons show cells containing increasing numbers of single-guide RNAs (hereafter, guides) as MOI increases. **B,** Histogram of the number of guides per cell, pooled across all six MOI levels (N=93 305 cells, mean=3.69 guides per cell). Vertical dashed lines show the median number of guides per cell. **C,** Histogram of cells per promoter-targeting guide, normalized to 100 000 total cells and pooled across all six MOI levels. **D,** Histogram of the number of guides per cell, split across increasing MOI levels. **E,** Heatmap of predicted transcript abundance changes with increasing number of guides per cell for 606 guide burden-responsive genes (F-test, BH-adjusted *p ≤* 10*^−^*^3^). **F,** Predicted transcript abundance changes of guide burden-responsive genes split into two clusters. Cluster 1 (left, 167 genes), cluster 2 (right, 320 genes). **G,** Gene ontology (GO) term analysis of cluster 1 (left) and cluster 2 (right). Benjamini-Hochberg (BH) adjusted p-values are shown by color.

To quantify the effect of guide burden on transcriptomic responses, we pooled cells across the six MOI conditions (88 287 cells with one or more guides) and analyzed responses using a modified version of Limma-voom [16] (Methods 1.10.2) with spline terms to model gene expression changes with increasing guide burden (Methods 1.8). We hypothesize that increasing guide burden may lead to transcriptional responses independent of the gene knockdown effects, because each guide corresponds to a lentivirus integrating into the genome [17]. For each cell, we modeled the effect of each perturbation, the number of guides detected in that cell (guide dosage), and cell quality covariates (fraction of ribosomal reads, fraction of mitochondrial reads, nuclear fraction of reads; Methods 1.6), as well as batch effects for each of the six MOI levels. We modeled the guide dosage as a cubic spline, allowing us to flexibly model the effects of increasing guide burden from zero to fifteen guides (Methods 1.8). This model allows us to find transcriptional responses due to increasing guide dosage, while controlling for the effect of genetic perturbations.

We predicted 606 genes to have a statistically significant guide burden effect (F-test, BH-adjusted p-value *≤* 10*^−^*^3^, Methods 1.8.2). Visualizing the predicted transcript abundance changes with increasing guide burden, we found that these genes monotonically increased or decreased with an increasing number of guides per cell (Figure 1E). After applying K-means to cluster the dynamics into three clusters, we focused on the two largest clusters to analyze the biological effects of increasing guide burden. In cluster 1 (167 genes), we found genes whose expression increased with guide burden, such as stress- and injury-associated markers *GDF15* [18], *TXNIP* [19], *FTH1*, *LGALS3*, and *AREG* (Figure 1F, left). Gene ontology (GO) analysis of this cluster identified enrichment for oxidative phosphorylation, mitochondrial ATP synthesis, respiratory electron transport chain, and cytoplasmic translation (Figure 1G, left). In contrast, cluster 2 (320 genes) contained genes whose expression decreased with guide burden, including proliferation- and cell-cycle-associated genes such as *MKI67*, *TOP2A*, *CENPF*, and *PCLAF* (Figure 1F, right). GO term analysis found strong enrichment for chromosome segregation, nuclear division, mitotic nuclear division, and sister chromatid segregation (Figure 1G, right). Of note, these responses are likely independent of p53 because DLD-1 cells are *TP53* mutants [20]. The spline model reveals that these gene programs alter progressively with guide burden. Together, these results suggest that while moderate guide numbers are tolerated, extreme guide overloading in high-MOI Perturb-seq designs eventually activates the integrated stress response [21] and suppresses proliferative cell-cycle progression [22, 23].

### PerturbMatch enables scalable analysis of guide multiplets

To choose a simple multiplet analysis strategy that scales to millions of cells, we evaluated a marginal modeling approach that (1) omits interaction effects between guides and (2) enables per-perturbation parallelization.

Since more than half of our analyzed cells were guide multiplets and our targeted library concentrated on only 45 promoters, we quantified the strength of interaction effects between different perturbations. If interaction effects could be ignored, this would greatly reduce the size of the model. To assess whether these terms could be omitted, we compared interactions between perturbations versus interactions involving technical covariates (nuclear fraction, ribosomal and mitochondrial read fractions, and total counts). To do so, we extended reluctant interaction modeling [24] to the multivariate setting, allowing us to rapidly screen 1485 interactions for effects across transcriptomic readouts (Methods 1.9). We found technical covariates to dominate the interaction score, with guide interaction effects far below that of technical covariates (Supplementary Figure 1A). The largest guide interaction effect found was CTNNB1 and MYC, but this effect was smaller than the interaction between MYC and the technical factors “total counts” and “fraction of mitochondrial reads” (Supplementary Figure 1B, C). Finally, we included the top five interaction effects into the model and compared the log fold changes with and without the interactions; we found that they were nearly identical (Pearson correlation = 0.9774, Supplementary Figure 1D). Overall, this analysis suggests that ignoring interaction effects is a valid strategy to simplify perturbation analysis.

To further scale the analysis, we evaluated fitting one differential expression model per genetic perturbation [16, 25–28] versus fitting one full model across perturbations [2, 14, 29]. If marginal modeling is comparable to full, then we would favor the marginal because it could be scaled by parallelizing fits across perturbations. In each marginal model, we assigned cells into a treatment and control group. For a given genetic perturbation, cells containing at least one gene-targeting guide were assigned to the treatment group, while cells carrying at least one non-targeting guide (NT) and no guides targeting that perturbation were designated as controls. Because NT guides are more abundant than any individual targeting guide, naive grouping would produce control cells with lower guide burden than treatment cells.

We developed PerturbMatch, which first stratifies cells by guide multiplet class (Figure 2A) and then applies propensity score matching within each class to balance technical covariates between treatment and control cells (Supplementary Figure 1E; Methods 1.10.1). Because the marginal approach allows the same multiply perturbed cell to be used in different treatment groups, the effective number of cells per guide can increase even if the actual number of cells remains constant. Indeed, normalizing to 10 000 total cells to compare across experiments, the effective number of cells per guide increased from 225 at baseline to 1600 as we increased MOI, a 7.1-fold gain (Supplementary Figure 1F). Across all levels of MOI, the inferred LFCs from the marginal model agreed with the full model (Figure 2B, Supplementary Figure 2A). Comparing LFCs inferred from marginal versus full model (fit with Limma-voom), we found the Pearson correlation decreased from 0.92 to 0.85 with increasing MOI levels (Figure 2C). The high Pearson correlation suggests that the marginal approach is comparable to the full model.

**Fig. 2.**
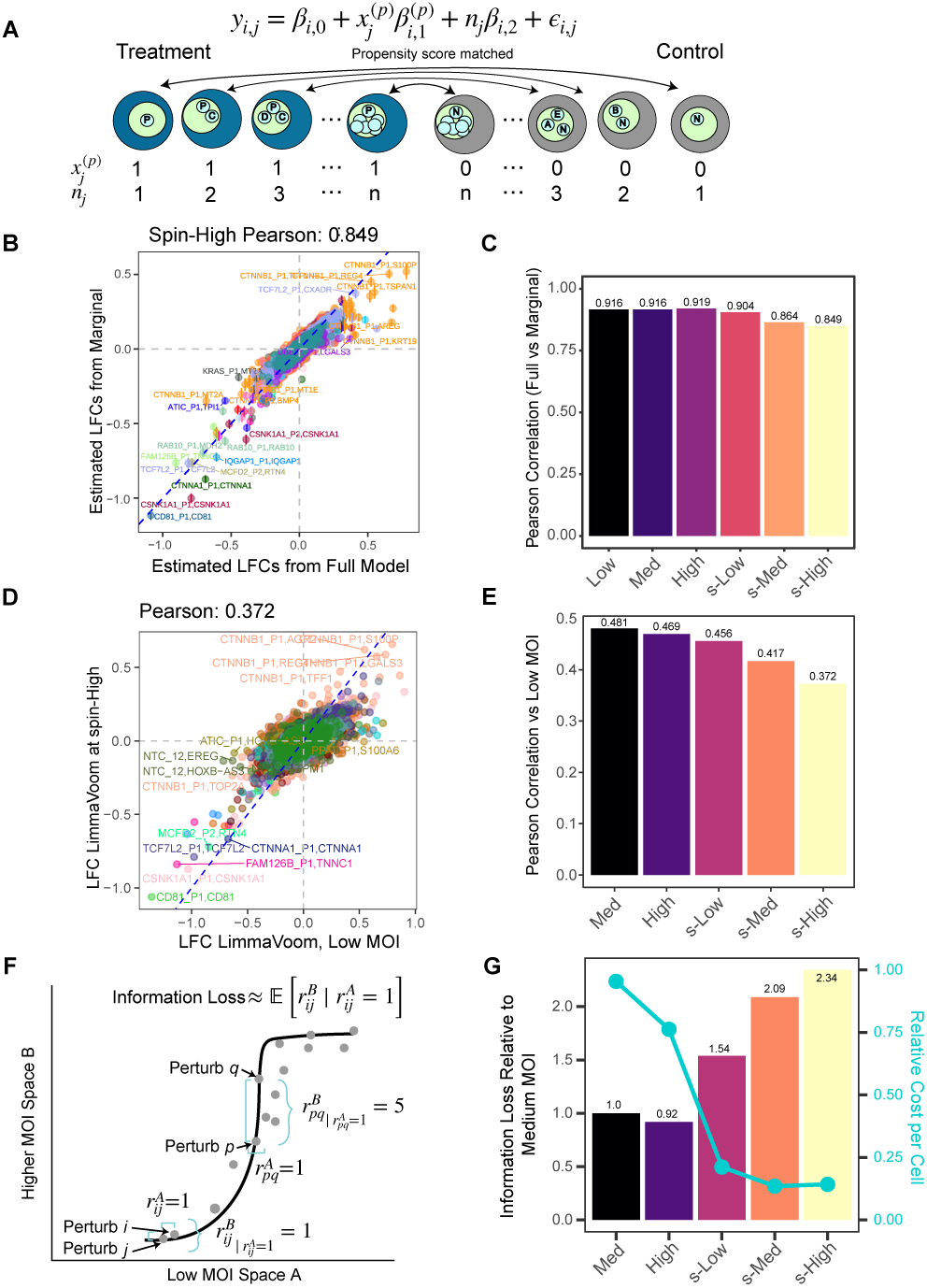
Analysis of guide multiplets quantifies the trade-off between information loss and cost savings. **A**, Illustration of PerturbMatch to balance covariates and model transcriptional response. Guides are shown as lettered circles inside cells. Treatment cells (*x^p^_j_* = 1) contain at least one guide *P* and control cells (*x^p^_j_* = 0) contain at least one NTC guide *N* and no *P* guide. Stratifying cells by the number of guides *n_j_*, we use propensity scores to select control cells with technical covariates similar to those of treatment (Methods 1.10.1). **B,** Scatterplot of LFC estimates from a full model versus a marginal model. Comparisons were performed at the highest MOI level. Pearson correlation is calculated across all perturbations and genes. Different colors indicate knockdowns of different gene targets. **C,** Barplot of Pearson correlations between LFC estimates from the full and marginal models across MOI levels. The prefix “s-” indicates spinfection. **D,** Scatterplot of LFC estimates in the low versus the spin-high MOI levels. Pearson correlation is calculated across all perturbations and genes. Different colors indicate knockdowns of different gene targets. **E,** Barplot of Pearson correlation summaries of low MOI LFCs versus the other five MOI levels. **F,** Illustration of neighbor ranks to calculate information loss (Methods 1.11). Perturbations *i* and *j* are nearest neighbors in both space A and B, therefore information is preserved for *i* and *j*. Perturbations *p* and *q* are nearest neighbors in space A, but not in B, therefore information found in A is lost in B for *p* and *q*. Information loss summarizes this rank displacement across all pairs of points. **G,** Barplot of normalized information loss across MOI levels, expressed as the fold change relative to the loss observed between low and medium MOI (Methods 1.11). The cyan line shows cost per cell for each condition, relative to low MOI.

We also compared Limma-voom with other differential expression methods. Of note, we compared estimates between marginal model Limma-voom and full model FR-Perturb [14], and found some mRNA abundances were downregulated in Limma-voom but not FR-Perturb (Supplementary Figure 2B). This likely reflects FR-Perturb having difficulty inferring LFCs that were not shared across perturbations, such as the downregulation of a target gene’s mRNA (Supplementary Figure 2C,D). We also compared Limma-voom against glmGamPoi [27], SCEPTRE [26], and ElasticNet [29], and found that they were mostly comparable with each other (Supplementary Figure 2E). Our marginal models took less than one minute to fit all 50 perturbations in parallel, while ElasticNet with five-fold cross-validation took up to three hours. For thousands of perturbations, the wall-time to fit marginal models would remain roughly unchanged (assuming parallel processing), while ElasticNet would likely increase to days. Favoring simplicity, speed, and scalability, we combined propensity scores with marginal Limma-voom (hereafter referred to as PerturbMatch) for all subsequent analyses.

### Quantifying the trade-off between information loss and cost savings in high-MOI designs

The increase in the effective number of cells at high MOI likely comes at the cost of reduced signal due to passenger guide and lentiviral effects. Indeed, global Pearson correlations across 3022 transcript LFCs and across the 45 perturbations between low MOI versus higher MOI levels progressively deteriorated from 0.48 to 0.37 (Figure 2D, E, Supplementary Figure 3A). We compared this information loss with the cost savings. We calculated the relative information loss at increasing MOI levels by comparing the rank of the nearest neighbors, which is a discrete analogue to copula variables [30] (Figure 2F, Methods 1.11). Specifically, for each perturbation *i*, we identified its nearest neighbor *j* in the reference experiment and then determined the rank of *j* among the neighbors of *i* in the query experiment using Euclidean distances between their transcriptome-wide logFC z-score vectors. If *i* and *j* are also nearest neighbors in the query experiment, then there is no information loss for *i* and *j*, but if the rank is greater than one, then there is loss. We averaged this quantity across all perturbations to determine the mean rank displacement. We call this quantity rank information loss or simply information loss. Using the loss from low to medium MOI as the baseline, we calculated the loss as fold change relative to baseline for each of the higher MOI levels. In this targeted Perturb-seq experiment, we can reduce cost per cell by 86% while maintaining information loss within two-fold of baseline (Figure 2G). Overall, our analysis systematically quantifies information versus cost trade-offs in high-MOI designs.

### Two- and three-guide multiplets best recover singlet perturbation responses

To evaluate the impact of guide multiplets (doublets, triplets, and higher-order multiplets) on information loss, we performed a computational titration by replacing singlets with guide multiplets while maintaining a constant number of cells. We used singlets from the lowest MOI (*N* = 15 598) to generate reference sets of differentially expressed genes (DEGs, absolute LFC *>* 0.1 and local false sign rate *<* 0.01; Methods 1.13). Using those DEGs as the reference ground truth across 45 perturbations, we calculated the mean precision, recall, and F-score at increasing fraction of guide multiplets. We also calculated cosine similarity of the z-scores of LFCs across the 3022 transcripts between the same perturbation at increasing fraction of guide multiplets. As a holdout baseline, we sampled 15 598 singlets in the higher MOIs (Figure 3A). This number was fixed while we replaced singlets with guide multiplets. Comparing reference with holdout, we calculated the recall, precision, F-score, and cosine similarity (*f* = 0, Figure 3B). We compared this baseline with the metrics calculated for six guide multiplet fractions (*f* = 0.1 to *f* = 1.0; Figure 3B). We found that mean precision was high (greater than 0.80) across all multiplet fractions and all multiplet types (Figure 3B). Guide doublets and triplets achieved higher mean recall than higher-order multiplets (Figure 3B). As the fraction of quadruplets (*n* = 4) or higher-order multiplets (*n ≥* 5) increased, recall, F-score, and cosine similarity declined rapidly (Figure 3B). In contrast, these metrics were more stable as the fraction of doublets or triplets increased. Perturbations with low baseline reproducibility—defined by low cosine similarity between holdout and reference singlets—also tended to have low cosine similarity in multiplet-containing query sets (Figure 3C). This suggests that perturbations with weak or absent transcriptomic phenotypes are difficult to reproduce at the available sample sizes. Across most multiplet fractions and multiplet types, cosine similarity of query versus reference correlated with baseline, with less concordance in higher-order multiplets (*n ≥* 4) (Supplementary Figure 3B). Overall, doublets and triplets best recapitulated the singlet perturbation responses, especially when mixed with singlets. Pure quadruplets and quintuplets had lower cosine similarity than baseline, likely due to noise from passenger guides. We hypothesize that this noise would be reduced in unbiased, large-scale screens, where most passenger guides would have negligible transcriptome effect.

**Fig. 3.**
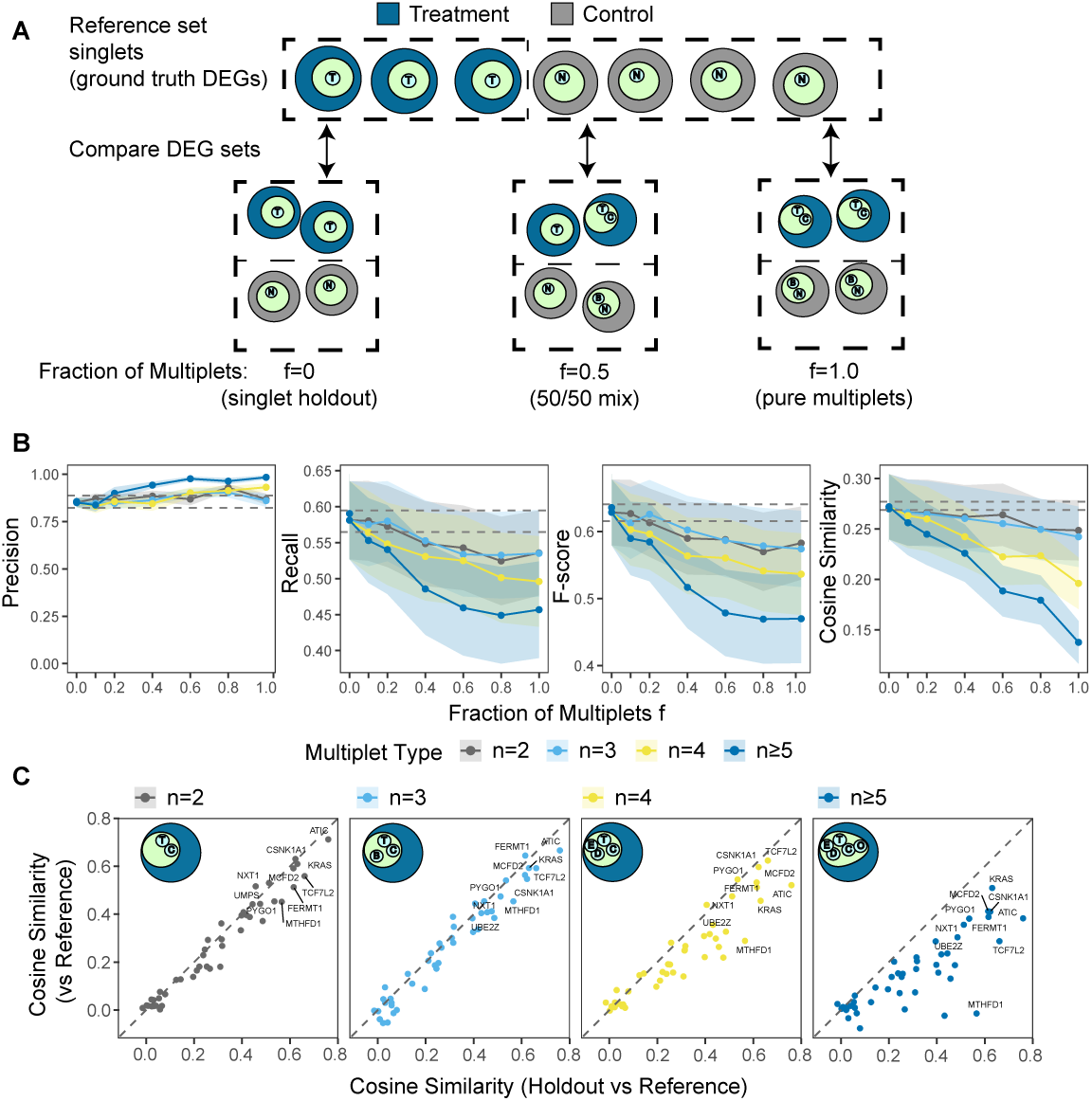
Two- and three-guide multiplets best recover singlet perturbation responses in computational titrations. **A**, Schematic of the computational titration framework using guide doublets as an example. Dark blue cells are treatment cells because they contain targeting guide *T*, and light grey cells are control cells because they contain a non-targeting control guide *N*. The reference set consists of low-MOI singlets (N=15 598 cells), which are used to define ground-truth DEGs. Each query set also contains 15 598 cells, but with varying fractions of guide multiplets. **B,** Mean precision, recall, F-score, and cosine similarity at increasing multiplet fractions. Each line represents a specific multiplet type. Ground-truth DEG sets are derived from reference singlets, with one DEG set per perturbation (45 perturbations total). Cosine similarity is calculated between transcriptome-wide LFC vectors from each query set and the reference. Shaded ribbons indicate *±*1 standard error across perturbations. Horizontal dashed lines represent the minimum and maximum metric values across seven random holdout-singlet subsets of 15 598 cells. **C,** Scatterplots comparing cosine similarity between holdout and reference singlets (x-axis) with cosine similarity between multiplet-based estimates and reference singlets (y-axis), shown separately for each multiplet type.

### High-MOI designs reduce cost while preserving perturbation signal

To evaluate the high-MOI approach in large-scale Perturb-seq screens, we performed three Perturb-seq screens targeting a quarter of the genome (5943 promoters targeting 4954 genes, hereafter called “5000-gene Perturb-seq”) at low, spinfection medium (hereafter, medium), and spinfection high (hereafter, high) MOI (Figure 4A). We obtained 1 869 846 cells after QC, of which 1 432 753 had at least one assigned guide. Among guide-positive cells, the mean guides per cell were 1.2, 3.7, and 6.7 in low, medium, and high MOI respectively (Figure 4B). Because an individual cell with multiple guides could be included in many marginal models, the medium MOI yielded three times more cells per guide than low MOI, and high MOI yielded five times more (Supplementary Figure 4A-C).

**Fig. 4.**
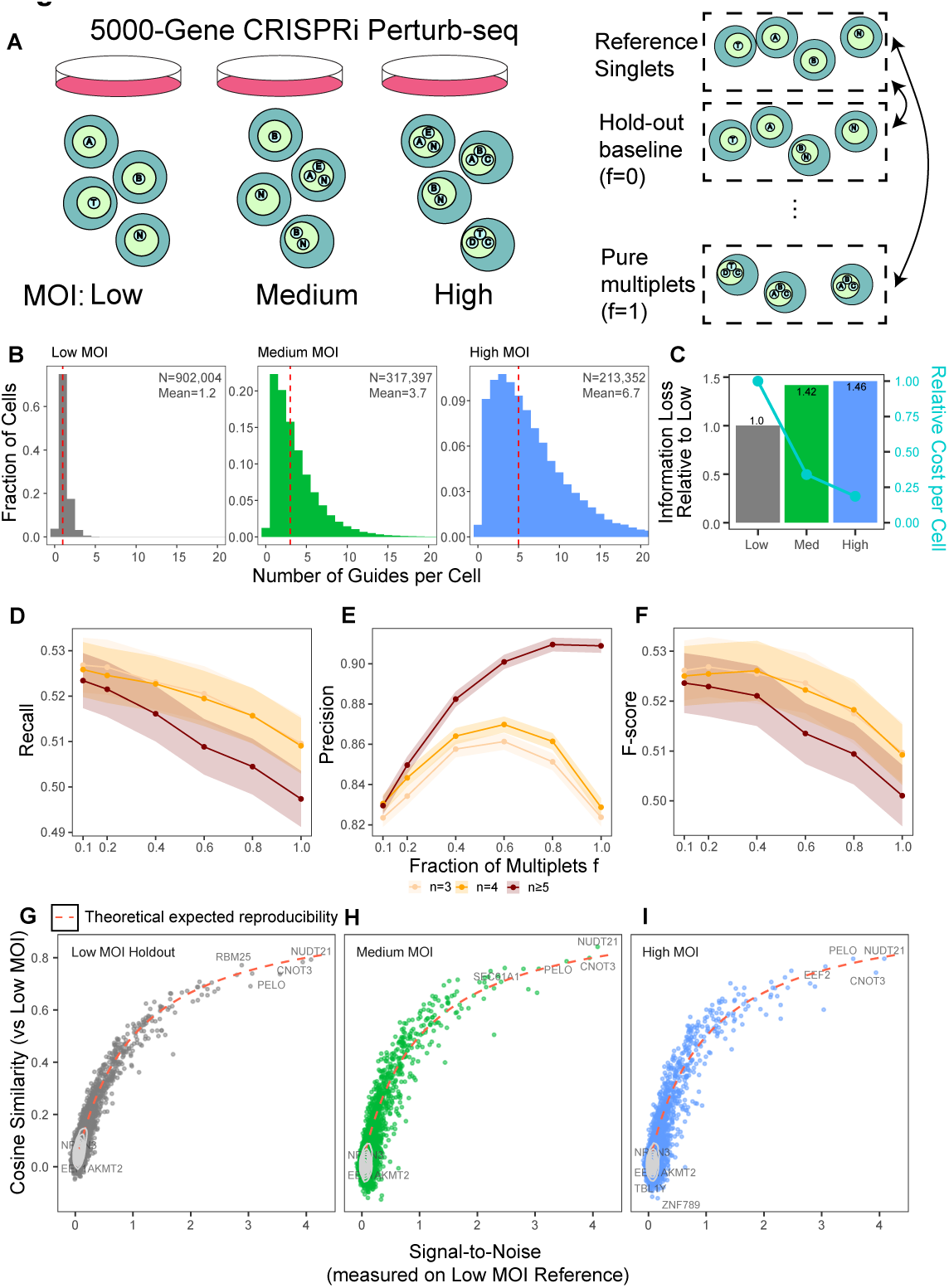
High-MOI designs reduce cost while preserving perturbation signal. **A**, Schematic of a quarter-genome Perturb-seq experiment targeting 5943 promoters across 4954 genes at three increasing MOI levels. **B,** Histogram of the number of guides per cell at the three MOI levels. **C,** Barplot of normalized information loss, expressed as fold change relative to the loss observed between low-MOI reference and low-MOI holdout (Methods 1.11). The cyan line shows per-cell costs relative to low MOI. **D,** Mean recall across 5943 perturbations at different multiplet fractions and multiplet types. Shaded ribbons indicate *±*1 standard error across perturbations. **E,** Same as **D**, but for precision. **F,** Same as **D**, but for F-score. **G-I,** Scatterplots of cosine similarity between each query set and the low-MOI reference, plotted against the signal-to-noise ratio (SNR) of each perturbation. Query sets are low-MOI holdout singlets (**G**), medium MOI (**H**), and high MOI (**I**). The dashed curve shows the theoretical reproducibility benchmark as a function of SNR (Methods 1.12).

We again quantified information loss with increasing MOI similar to our 35-gene Perturb-seq data. We split the low MOI cells in half to create a reference and a holdout set. We used the information loss between reference and holdout to normalize information loss at higher MOI. We found that the losses between the reference and medium or high MOI were within two-fold of the baseline (Figure 4C). Of note, this loss was smaller than the high MOI levels of the 35-gene Perturb-seq experiment. This smaller loss likely reflects the unbiased guide RNA library compared to the 35-gene Perturb-seq. With large-scale unbiased libraries, most passenger guides in multiplets would contribute little or no transcriptomic response. This relatively small loss comes with large cost savings: medium MOI reduced cost by 66% and high MOI by 81% per cell, relative to low MOI (Figure 4C). Therefore, compared with our previous 35-gene targeted screen, our 5000-gene screen shows less information loss, suggesting that the high-MOI strategy better preserves perturbation signal, consistent with reduced passenger-guide interference in large libraries.

Similar to our targeted screen, we then dissected the contributions of different guide multiplet types. We again fixed the total number of cells in a holdout set and performed a computational titration to gradually increase multiplet fraction. We used DEGs defined from 676 146 singlets in low MOI as reference to estimate recall and precision of query cells. Because there were relatively few singlets at medium and high MOI (N=90 217), we also included doublets (N=85 919) in our holdout set to increase the number of DEGs called. Guide multiplets from low MOI were omitted to minimize shared batch effects. In total, we had 176 136 holdout cells for the titration analysis.

Across all types of guide multiplets tested, mean recall (across 5943 promoters) deteriorated slightly as a function of multiplet fraction (from 0.525 to 0.5, Figure 4D). Precision was high (*>* 0.8) in all three guide multiplet classes (Figure 4E). For guide triplets and quadruplets (*n* = 3 and *n* = 4), precision peaked between 0.6 and 0.8 multiplet fraction (Figure 4E). In contrast to our targeted screen, pure quintuplets or greater (*n ≥* 5) performed similarly to triplets and quadruplets, suggesting that noise from passenger guides in large unbiased screens is smaller than in the targeted screens (Figure 4F). We compared cosine similarities in holdout versus reference for each of the 5943 perturbations and asked how they correlated with similarities in medium and high MOI versus reference (Supplementary Figure 4D,E). We found high concordance between the two cosine similarities, with most cosine similarities near zero.

We reasoned that since targeting randomly selected genes usually leads to insignificant transcriptional responses, many cosine similarities between replicates should be close to zero. Cosine similarities that were high in both query and baseline had large changes in the transcriptome, quantified by the root mean square of the standardized LFCs (RMS, Supplementary Figure 4D,E, Methods 1.12).

### Response magnitude bounds expected signal recovery

We derived a theoretical framework to model the relationship between the magnitude of the transcriptomic response, quantified as RMS, and its expected reproducibility, measured by cosine similarity. Because the z-scored LFCs are standardized to a unit noise floor (i.e. variance equals one), the biological signal power is numerically equivalent to the signal-to-noise ratio (SNR = RMS^2^ *−* 1). Under an additive noise model, where biological signal is shared but noise is approximately orthogonal, the expected cosine similarity *ρ* would be the proportion of signal variance to the total variance: 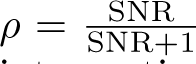. This relationship provides an expected reproducibility benchmark when interpreting cosine similarity (Methods 1.12).

For each query set (low holdout, medium, high MOI) we summarized the 5943 perturbation responses by their SNR and cosine similarity with the reference. The empirical relationship between SNR and similarity agreed well with our theoretical model (Figure 4G-I); strong perturbation responses yielded high similarity, while weak ones showed a predictable decline in concordance. Interestingly, most genes in our unbiased 5000-gene Perturb-seq screen responded weakly, limiting their maximum reproducibility. Finally, the close agreement between our empirical cosine similarities and the theoretical expected reproducibility demonstrates that the medium and high MOI scenarios can recover the reference signal at lower costs.

### Incorporating guide multiplets improves statistical power and signal recovery in genome-wide Perturb-seq

To apply our joint analysis to existing genome-scale data, we reanalyzed the Replogle 9868-perturbation screen in K562 cells to evaluate statistical power with and without guide multiplets. We split the 2 392 277 singlets (*n* = 2 guides per cell due to their dual targeting strategy) into a reference (80%, 1 913 822) and holdout set (20%, 478 455), and used 461 146 guide doublets (*n* = 4) and 81 649 guide triplets (*n* = 6) as multiplets (Figure 5A, B, C). Cosine similarity between the reference and singlet-only query was similar to that between the reference and multiplet-only query (Figure 5D). Notably, the joint query of singlet holdout plus multiplets had higher cosine similarity to the reference than the singlet-only query (Figure 5E).

**Fig. 5.**
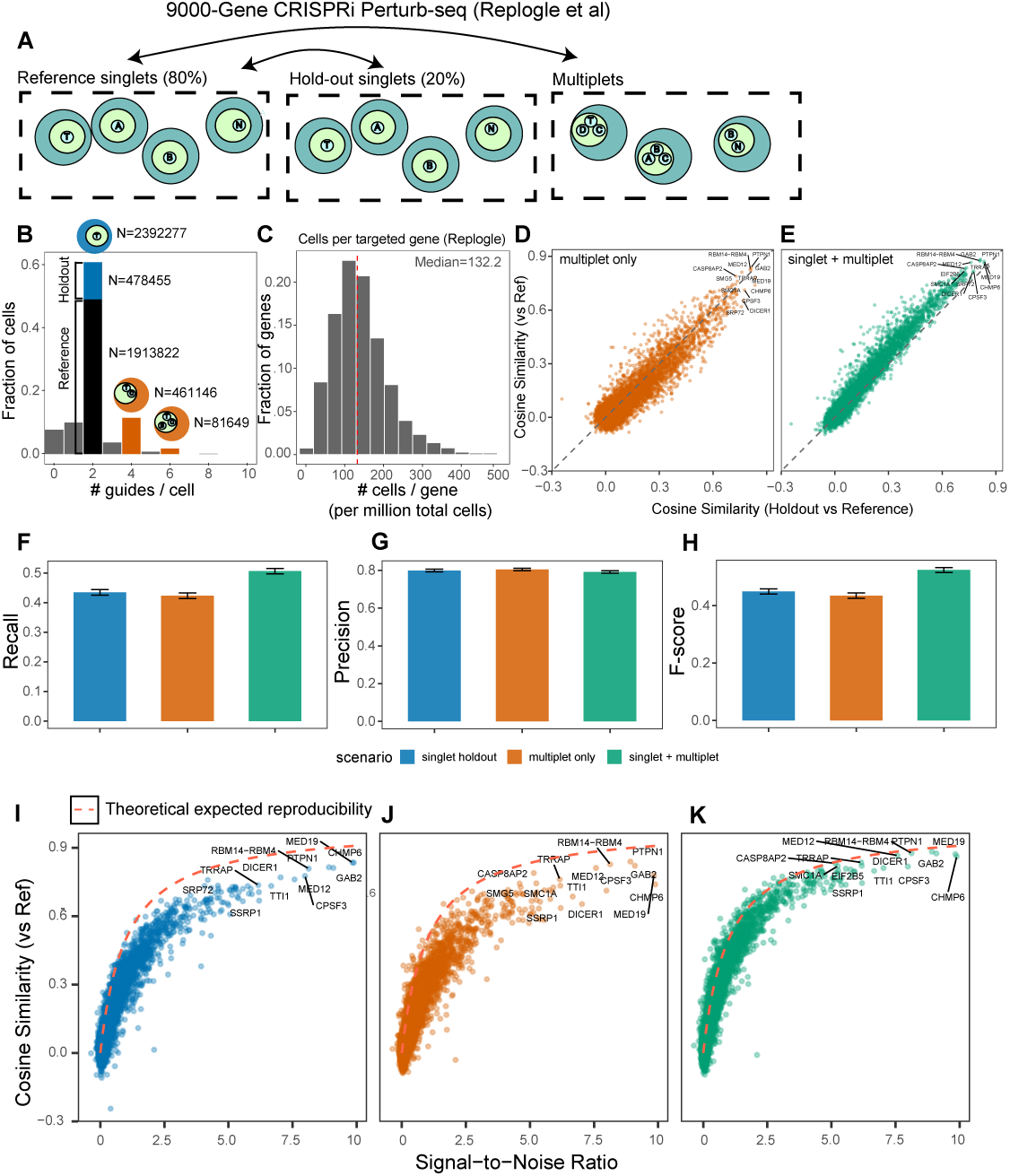
Incorporating guide multiplets improves statistical power and signal recovery in genome-wide Perturb-seq. **A**, Schematic of the power analysis in the Replogle et al. Perturb-seq dataset. Singlets are split into 80% reference and 20% holdout sets. Guide multiplets, defined as cells containing two or three genetic perturbations, are kept as an additional query set. The analysis compares three query scenarios against the reference: holdout singlets, multiplets only, and holdout singlets plus multiplets. **B,** Histogram of the number of guides per cell from the dual-guide sgRNA strategy. Singlets, doublets, and triplets are located at two, four, and six guides per cell, respectively. **C,** Histogram of the number of cells per targeted gene, normalized to one million total cells. Vertical dashed line indicates median at 132.2 cells per gene. **D,** Cosine similarity between the multiplet-only query and reference, plotted against cosine similarity between holdout singlets and reference. Each point is one perturbation. **E,** Same as **D**, but for the joint query of holdout singlets plus multiplets. **F-H,** Mean recall, precision, and F-score across the three query scenarios: holdout singlets, multiplets only, and holdout singlets plus multiplets. Error bars indicate *±*2 standard errors across perturbations. **I-K,** Cosine similarity between each query scenario and reference, plotted against the signal-to-noise ratio (SNR) of the transcriptome-wide response calculated from the reference. Panels show holdout singlets (**I**), multiplets only (**J**), and holdout singlets plus multiplets (**K**). The dashed curve shows the theoretical reproducibility benchmark as a function of SNR (Methods 1.12).

To quantify this further, we calculated mean recall and precision using DEGs defined from reference as ground truth. The multiplet-only scenario yielded mean recall and precision comparable to those of the 20% singlet holdout, whereas recall improved significantly in the joint scenario (Figure 5F,G). The joint scenario also yielded a higher mean F-score than either the singlet-only or the multiplet-only scenario (Figure 5H). At the perturbation level, recall and F-score were generally higher in the joint scenario than in the singlet-holdout scenario, whereas precision was similar between the two (Supplementary Figure 5A-C).

We evaluated the relationship between transcriptional response magnitude and reference concordance using the same framework as in the 5000-gene Perturb-seq analysis. We found that singlet- and multiplet-only scenarios consistently fell below the theoretical expected reproducibility (Figure 5I, J), suggesting that those scenarios were underpowered. Remarkably, the joint scenario converged upon the theoretical expected reproducibility, improving signal recovery globally across perturbations (Figure 5K). This improvement is likely due to an increase in the number of cells per perturbation, which boosts statistical power (Supplementary Figure 5D). Overall, we demonstrate that joint analysis with multiply perturbed cells even in low MOI settings can improve statistical power.

## Discussion

Loading multiple guides into each cell has emerged as a practical strategy for reducing the cost of targeted genetic [14] and enhancer Perturb-seq screens [6, 12]. However, adopting and deploying cost-efficient Perturb-seq at the scale demanded by foundation models requires clarifying lingering uncertainties regarding biological viability, data quality, and computational tractability.

Here we show—using targeted Perturb-seq, quarter-genome Perturb-seq, and genome-wide Perturb-seq—that guide multiplets can be incorporated into the analysis while preserving perturbation signal. For high-MOI designs, our results suggest several practical guidelines. First, moderate guide overloading can preserve high-quality perturbation signal, whereas extreme guide burden induces stress and suppresses cell-cycle progression. Second, cells carrying two or three guides recover perturbation responses better than higher-order multiplets. Third, perturbation-specific marginal models provide a simple and scalable strategy for analyzing guide multiplets across millions of cells. Overall, we recommend a Perturb-seq design that includes singlets, doublets, and triplets to improve cost efficiency while preserving signal recovery. We believe that this simplified experimental and analysis framework may make large-scale Perturb-seq experiments more practical for “lab-in-the-loop” paradigms [31], where cheap, easy, and fast iterations enable rapid hypothesis testing.

This work also introduces experimental, computational, and theoretical frameworks for interpreting perturbation screens more generally. Experimentally, we demonstrate that replicates—currently underutilized in large-scale Perturb-seq—are critical for separating reproducible perturbation responses from measurement noise. Computationally, our information-loss analysis provides a quantitative metric for comparing relative information content across experimental designs or measurement modalities. Theoretically, our model predicts expected reproducibility from the magnitude of the perturbation response. These frameworks may help guide iterative screen design [32], multimodal perturbation analysis [33, 34], and prediction of responses to unseen perturbations [31].

## Supplementary Figures

**Supplementary Figure 1.**
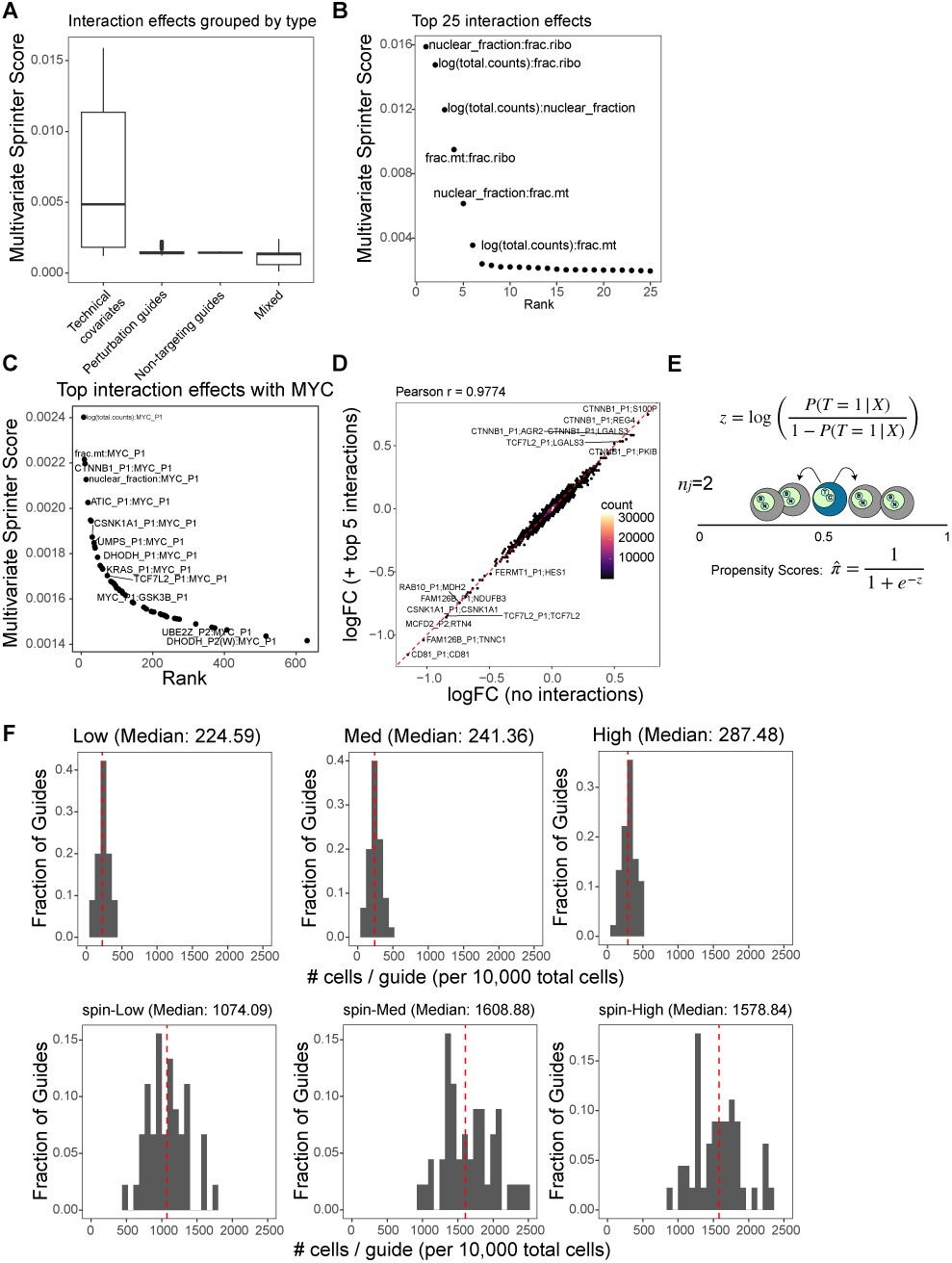
Interaction screening supports simplified marginal modeling of guide multiplets. **A**, Boxplot of potential interaction effects quantified by the multivariate sprinter score (Methods 1.9). Boxes show four categories of interactions: technical covariate interactions (total counts, mitochondrial read fraction, ribosomal read fraction, and nuclear fraction), perturbation-perturbation interactions, NTC-NTC interactions, and mixed interactions between technical covariates, perturbations, or NTCs. **B,** Top 25 two-way interaction effects ordered by the multivariate sprinter score. Labels indicate the two interacting covariates. **C,** Top two-way interaction effects involving the MYC perturbation. **D,** Comparison of LFC estimates from models fit with and without the top five interaction terms across 50 perturbations and 3022 genes. **E,** Schematic of the PerturbMatch propensity score matching procedure (Methods 1.10.1). Within each guide-count stratum *n_j_* (doublets, *n_j_* = 2, shown), we fit a weighted logistic regression model to classify treatment and candidate control cells using only technical covariates, with weights used to balance treatment and control class sizes. For *N* treatment cells, we match *K × N* control cells (*K* = 2 shown; *K* = 5 used in practice). **F,** Histogram of the number of cells per guide across the six MOI levels, normalized to 10 000 total cells to enable direct comparison across experiments.

**Supplementary Figure 2.**
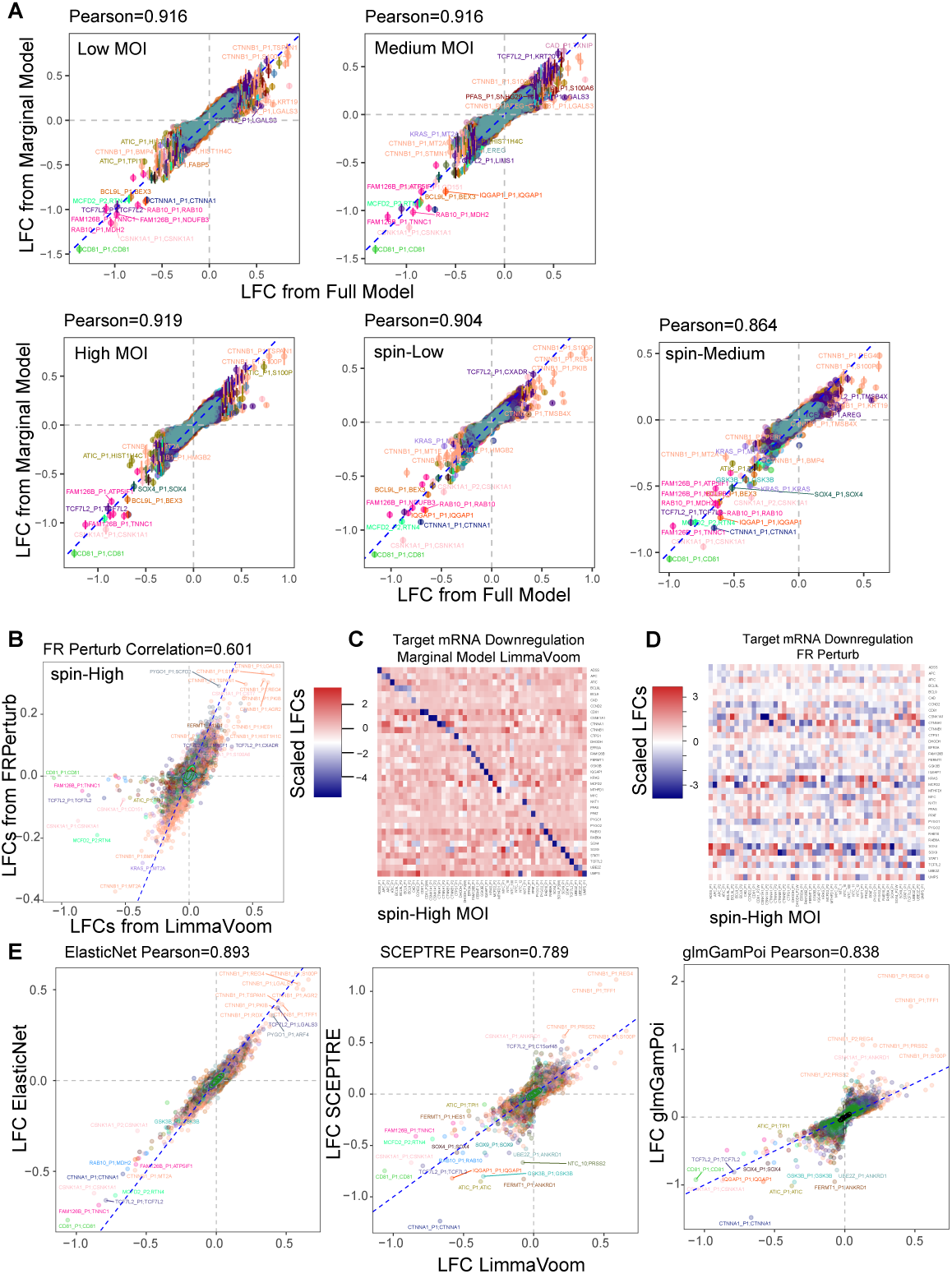
Model comparisons support marginal modeling for guide multiplet analysis. **A**, Scatterplots of LFC estimates from the full model versus the marginal model across MOI levels. Pearson correlations are calculated across all perturbations and genes. Colors indicate knockdowns of different gene targets. **B,** Scatterplot of LFC estimates from Limma-voom versus FR-Perturb [14]. Labels indicate the genetic perturbation followed by the measured mRNA, formatted as “perturbation;mRNA”. For example, “CD81 P1;CD81” indicates the effect of the CD81 P1 perturbation on CD81 mRNA abundance. Colors indicate genetic perturbations. **C,** Heatmap of target mRNA abundance changes after genetic perturbation, with LFC estimates from Limma-voom. Diagonal patterns indicate downregulation of the targeted mRNA after knockdown. LFCs are calculated for the highest-MOI condition (spin-High). **D,** Same as **C**, but using LFC estimates from FR-Perturb. **E,** Scatterplots of LFC estimates from ElasticNet, SCEPTRE, and glmGamPoi versus Limma-voom. ElasticNet is a full model, whereas SCEPTRE, glmGamPoi, and Limma-voom are marginal models.

**Supplementary Figure 3.**
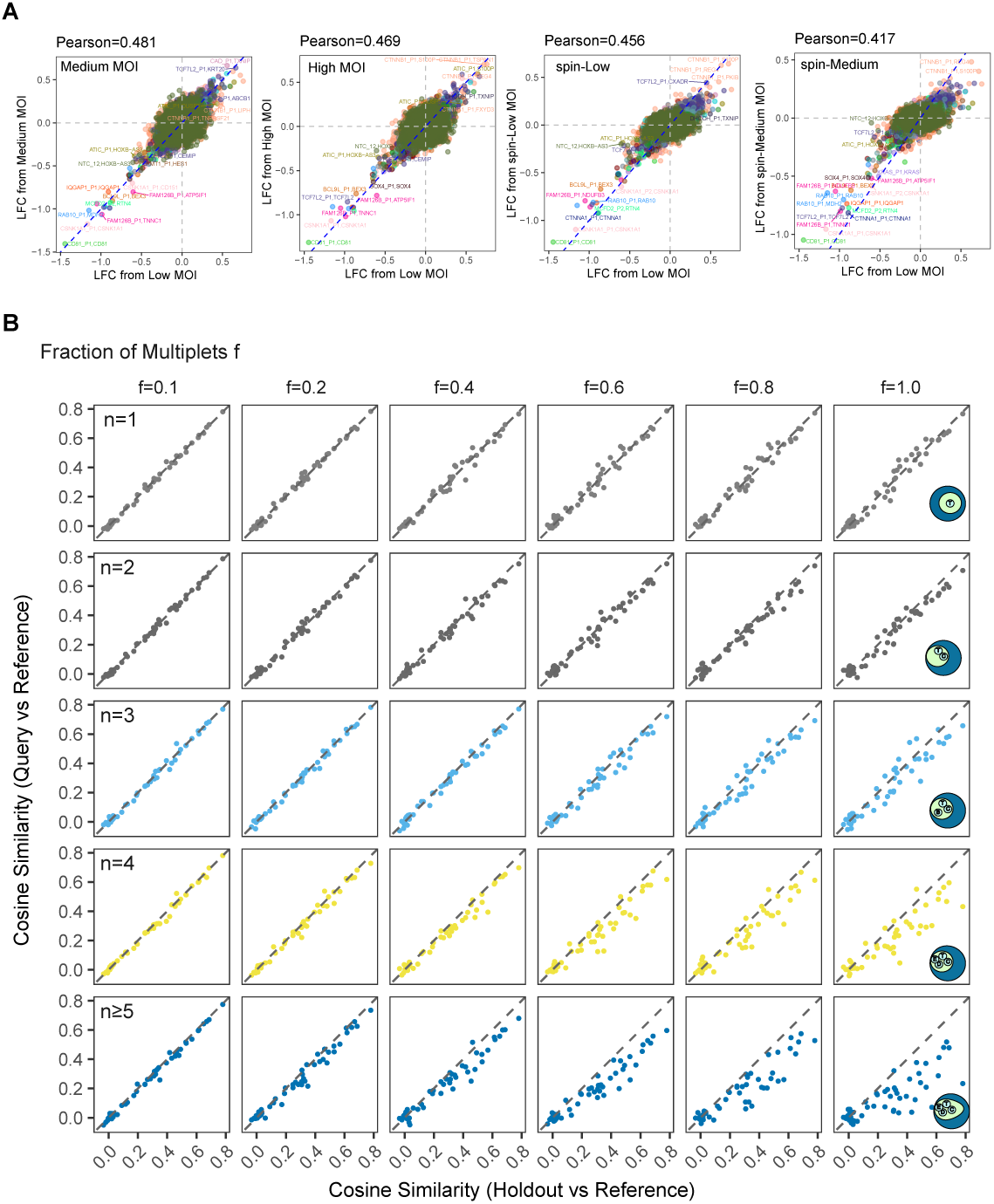
In targeted Perturb-seq, higher MOI and higher-order guide multiplets reduce signal recovery. **A**, Scatterplots of LFC estimates from low MOI versus higher MOI levels. Pearson correlations are calculated across all perturbations and genes. Colors indicate knockdowns of different gene targets. **B,** Scatterplots of cosine similarity between holdout and reference singlets (x-axis) versus cosine similarity between query and reference singlets (y-axis). Each query set is constructed by replacing increasing fractions of holdout singlets with multiplets. Each point represents one perturbation-level LFC profile. Columns show increasing multiplet fractions, and rows show different multiplet classes. Singlets (*n* = 1) classes are created by replacing a given fraction of holdout singlets with new singlets that are not present in either reference or the original holdout set.

**Supplementary Figure 4.**
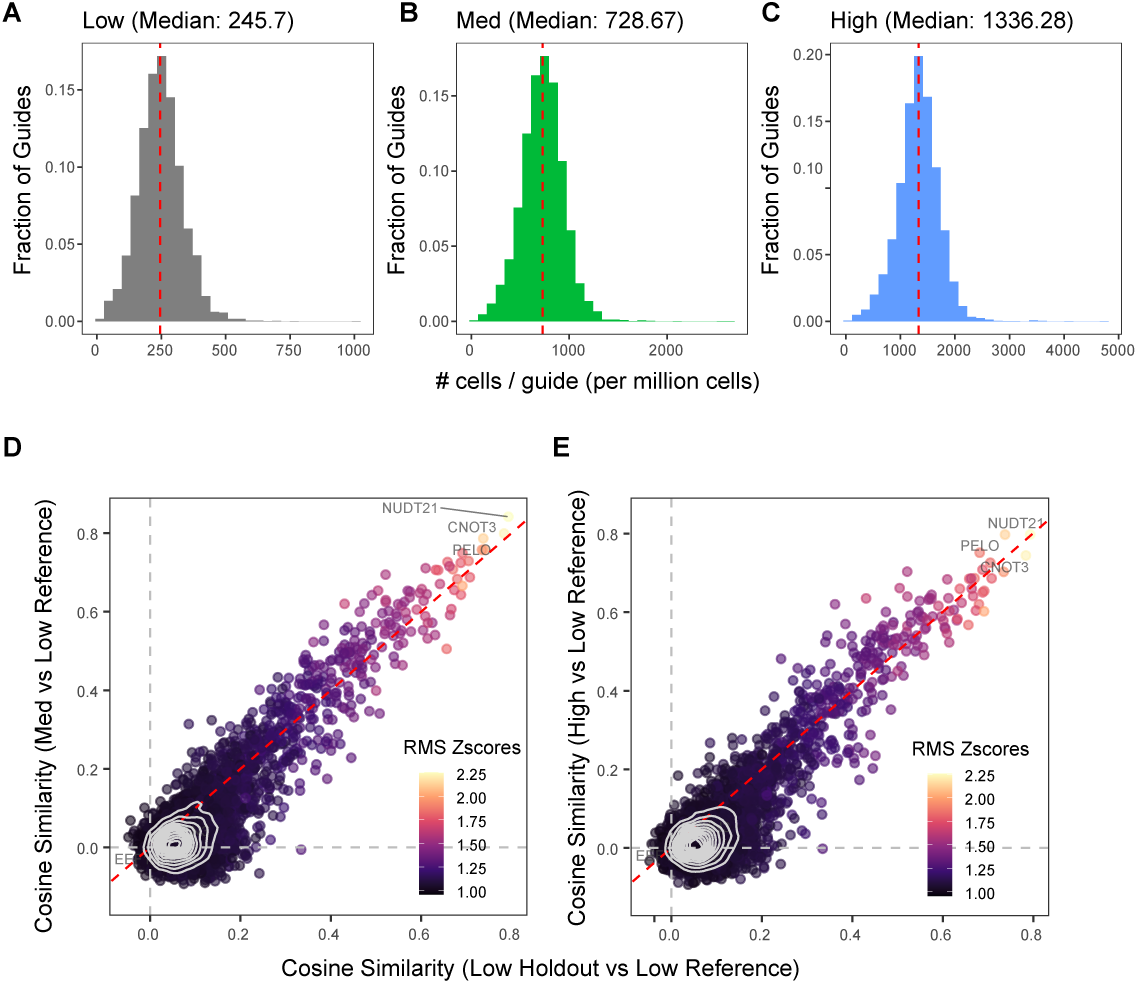
In large-scale, unbiased Perturb-seq, higher MOI increases the effective number of cells per guide and preserves perturbation signal. A-C, Histograms of the number of cells per guide across low (**A**), medium (**B**), and high (**C**) MOI levels, normalized to one million total cells. **D-E,** Cosine similarity between each query set and the low-MOI reference, plotted against cosine similarity between the low-MOI holdout and low-MOI reference. Query sets are medium MOI (**D**) and high MOI (**E**). Points represent perturbation-level LFC profiles and are colored by the root-mean-square of standardized LFCs (RMS Z-score), which is a measure of the transcriptome-wide impact of a perturbation.

**Supplementary Figure 5.**
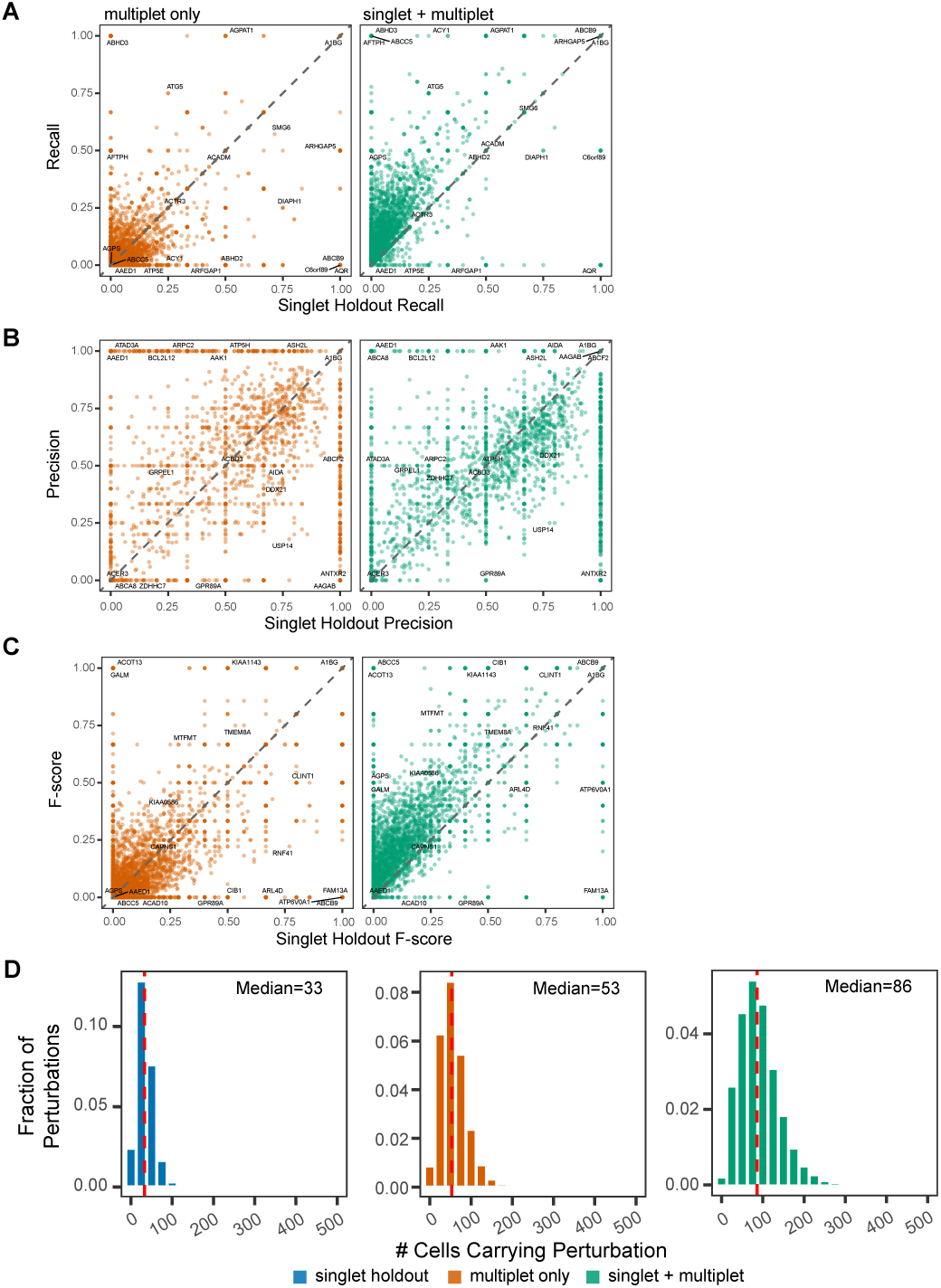
Joint singlet-multiplet analysis improves DEG recovery over singlet holdouts. A-C, Recall (**A**), precision (**B**), and F-score (**C**) for query sets plotted against the corresponding singlet-holdout metric. Ground-truth DEGs are calculated from the reference set of 80% of singlets. Left panels show metrics for DEGs derived from multiplets only. Right panels show metrics for DEGs derived from singlets and multiplets jointly. Each point represents one perturbation. Dashed lines indicate equal performance between the query set and singlet holdout. **D,** Histograms of the number of cells carrying at least one guide for each gene perturbation across the three query scenarios: singlet holdout, multiplet only, and singlet plus multiplet. Red dashed vertical lines indicate medians.

## 1 Methods

### 1.1 Cell lines

The DLD-1 colorectal cancer cell line and 293T lentiviral producer cells were obtained from the American Type Culture Collection (ATCC). DLD-1 cells were cultured in RPMI-1640 medium supplemented with 10% fetal bovine serum (FBS) and 2 mM L-glutamine. 293T cells were cultured in Dulbecco’s Modified Eagle Medium (DMEM) supplemented with 10% FBS, 100 µM non-essential amino acids (NEAA), and 2 mM L-glutamine. Both cell lines were maintained at an appropriate density by passaging every 2-3 days. Prior to experimentation, the identity of both cell lines was confirmed using short tandem repeat (STR) profiling, and they were routinely tested for mycoplasma contamination using commercially available kits to ensure they were mycoplasma-free.

### 1.2 sgRNA library design and construction

We designed two different sgRNA libraries to accommodate two different purposes. The sgRNA oligos were synthesized by GenScript and cloned into a CROP-seq-derived lentiviral vector [5]. The guides were designed to target the +/- 200 bp region of the promoter. For genes with multiple annotated promoters, we included all promoters in the library design. The four highest-scoring guides were selected to target each promoter, based on the on-target and off-target scores. The genes involved in the targeted library were chosen based on known biology of DLD-1 cells. For the unbiased screen, the 4954 genes were chosen randomly to resemble the distribution of genes in genome-wide screens.

### 1.3 Lentivirus packaging

One day before transfection, the cell culture dish was treated with 5 mL of 1% gelatin in PBS, incubated for 10 min, then aspirated. 3.5*×*10^6^ 293T cells were seeded into each 10 cm dish to reach 80% confluence. On the day of transfection, 20 µL Lipofectamine 2000 was added to 480 µL OptiMEM. In a separate tube, the sgRNA-library transfer plasmid, pDelta8.9 packaging plasmid, and pVSV-G envelope plasmid were mixed in 500 µL of Opti-MEM at 5, 16, and 1 µg, respectively. Both mixes were combined and incubated for 20 min before adding to the dish. The dish was incubated at 37°C for 6 h and then 10 mL of complete medium was added. Virus was then harvested 48 h after transfection. Briefly, all supernatants were harvested and filtered through a 0.45 µm filter bottle. The virus was concentrated using the Lenti-X concentrator (Takara Bio) following the manufacturer’s instructions and resuspended in 1 mL 1% BSA in PBS per dish. The concentrated virus was divided into 200 µL aliquots and stored at -80 °C until infection. The virus titer was determined by running a serial-dilution transduction assay in DLD-1 cells followed by puromycin selection and CellTiter-Glo assay.

### 1.4 Perturb-seq experiment

DLD-1 cells were engineered with the ultra-tight CRISPRi system with the DD tagging using lentivirus infection with Blasticidin selection [35]. The gene knockdown efficiency was validated by CD81 flow cytometry. For low MOI infections, we directly added empirically titrated virus volumes to a T175 flask with 3.68*×*10^6^ cells seeded. For high-MOI infections, spinfection was performed in 24-well plates with 8 µg/mL polybrene by centrifugation at 3,100 rpm for 45 min at room temperature. In the 35-gene Perturb-seq experiment, we performed three MOI levels using the direct method and three MOI levels using spinfection. In the 5000-gene Perturb-seq, we performed the low MOI level with the direct method, and medium and high MOI using spinfection. Complete medium was added to each well 5 hours after the infection. For both conditions, the cells were selected with 2 µg/mL puromycin for 7 days before freezing for further analysis. For Perturb-seq experiments, frozen cells were thawed and recovered for 2 days before adding 250 ng/mL doxycycline and 1 µM Shield1 ligand (Takara Bio) to induce the expression of Cas9. Cells were harvested 5 days after the induction, hashed with the MULTI-seq cholesterol-modified oligos (CMO) [36] and then loaded on the 10x Genomics Chromium X instrument with the 3’ GEM-X V4 kit. We overloaded the channel so that about 50 000 to 60 000 cells were expected to be recovered from each channel. The library construction was largely consistent with our previous paper [13], except that hashing libraries were constructed following the MULTI-seq protocol. The GEX, sgRNA, and CMO hashing libraries were then sequenced with Illumina NextSeq 2000 or NovaSeq-X platforms targeting sequencing depths of 20,000, 2,000, and 2,000 reads per cell, respectively.

### 1.5 Perturb-seq mapping and processing

Raw sequencing data were processed using the Single Cell PerturbSeq pipeline, which automates the integration of sample metadata and the generation of standardized samplesheets for local or cloud-based analysis. Specifically, the data were processed via the Cumulus framework [37]. Initial sequence alignment, filtering, and transcript quantification were performed using the Cumulus cellranger workflow for mapping against the GRCh38-2020-A reference genome. Following alignment, cellular barcodes were assigned to individual experimental conditions and specific guide RNAs (gRNAs) using the Cumulus demultiplexing module. This dual-indexing strategy ensured accurate cell-to-sample hashing and precise perturbation assignment for downstream single-cell analysis.

### 1.6 Adaptive threshold filtering for quality control of cells

To filter features, we used binomial deviance [38] to calculate highly variable genes. For the 5000-gene and Replogle differential expression analyses, we used top 3000 highly variable genes. For the 35-gene analysis, we used the top 3000 plus the target genes themselves, resulting in a total of 3022 genes.

To filter cells, we used classic filtering thresholds to initialize outlier cells, then refined the thresholds using a Gaussian mixture model (GMM).

First, per-cell technical metrics were computed: total UMI counts, mitochondrial transcript fraction (fraction of reads mapping to MT-genes), and ribosomal transcript fraction (fraction of reads mapping to RPS/RPL genes). Nuclear fraction—the proportion of reads deriving from unspliced (intronic) transcripts—was estimated independently from aligned BAM files using DropletQC [39]. Outlier cells were identified for each metric using median absolute deviation (MAD)-based thresholds (5 MADs from the median on log-transformed values) via scuttle::isOutlier [40]. Cells exceeding any individual threshold were initialized as low quality.

Next, to refine the QC thresholds, we used a three-component multivariate Gaussian mixture model to fit a joint distribution of the cells in technical covariate space. The three components were initialized using cells identified by MAD-based thresholds: “damaged” (high nuclear fraction outliers), “empty” (low nuclear fraction outliers), and “good” (cells passing all thresholds). The EM algorithm was run for 20 iterations. Cells assigned to the damaged or empty components with posterior probability *>* 0.5 were removed from downstream analysis.

### 1.7 Assigning guides to cells using fishash

We used the R package fishash (version 0.4.2) [15] to assign guides to cells, using an adjusted *p*-value (padj) cutoff of 0.01. Briefly, fishash treats the guide RNA UMI count matrix as a contingency table and identifies associations between cell and guide barcodes using a one-sided Fisher’s Exact Test. To address hidden confounding from ambient RNA and barcode swapping, which can lead to false associations via Simpson’s paradox, the method iteratively estimates signal and noise components to re-compute the test statistics. We also applied fishash to 9000-gene Perturb-seq data from Replogle et al. to estimate multiplets.

### 1.8 Quantifying the effect of guide burden on gene expression response

#### 1.8.1 Spline model to characterize gene expression dynamics as a function of guide burden

To characterize guide-burden effects, we used a full model that allowed gene expression to change flexibly as a function of the number of guides detected in each cell. We call this a full model, as opposed to a marginal model, because the full model uses all analyzed cells across all perturbations together in one fit, while marginal models subset cells and fit each perturbation separately. For each gene *g*, we fit a global linear model across cells using a Limma-voom framework. Let *Y_gi_* denote the observed count for gene *g* in cell *i*, and let *y_gi_* denote the corresponding voom-transformed log-expression value. The model coefficients were estimated separately for each gene using a shared cell-level design matrix:

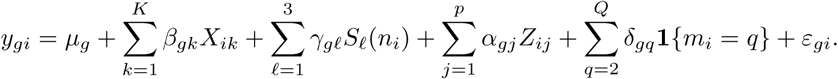

Here, *X_ik_* is the perturbation indicator for perturbation *k*, and *β_gk_* is the log-fold-change coefficient for gene *g* associated with perturbation *k*, conditional on guide burden, technical covariates, and MOI-level batch effects. In the pilot data, *K* = 50, representing 45 promoter perturbations plus 5 non-targeting control (NTC) guides.

The guide-burden effect is modeled as a function of a natural cubic spline in *n_i_*, the number of guides detected in cell *i*. Specifically, *S*_1_(*n_i_*)*, S*_2_(*n_i_*)*, S*_3_(*n_i_*) are the three natural cubic spline basis functions generated with three degrees of freedom, and *γ_gℓ_* is the coefficient for spline basis function *ℓ* in gene *g*. We used the R package splines to create a three-column natural cubic spline basis, implemented as `splines::ns(nguides.fishash, df = 3)`.

The term 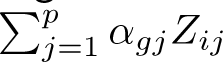 represents the technical covariates. In the spline fit, these covariates were nuclear fraction, mitochondrial fraction, and ribosomal fraction (*p* = 3). The variable *m_i_* ∈ {1*, . . ., Q*} indexes the MOI level for cell *i*, and *Q* is the total number of MOI levels. Because cells were pooled across six MOI levels in the pilot data, *Q* = 6. The term **1**{*m_i_* = *q*} is the indicator that cell *i* belongs to MOI level *q*, with *q* = 1 absorbed into the intercept; *δ_gq_* is the corresponding MOI-level intercept shift for gene *g*. *µ_g_* is the intercept for gene *g* at the reference MOI level (we explicitly set this to the low MOI), and *ε_gi_* is the residual error. This model assumes that perturbation contributions within the same cell are additive and that there are no perturbation-by-perturbation interactions. We discuss potential interaction effects further in Methods 1.9.

We used Limma-voom with a small adaptation to handle lowly expressed genes (Methods 1.10.2).

#### 1.8.2 Downstream analysis to identify and characterize genes responsive to guide burden effects

After fitting the full model, we performed a downstream analysis to identify genes whose transcript abundance changed significantly with guide burden and to visualize the fitted guide-burden trajectories. For visualization, the prediction grid was defined as a set of reference guide-count values *n*^∗^ spanning the interval from 1 to 15 guides per cell. We constructed a prediction design matrix with one row for each value of *n*^∗^, using the same natural-spline basis functions *S_ℓ_*(·) as in the fitted model.

Concretely, for a reference prediction profile evaluated at *n*^∗^ guides per cell, the design row can be written as:

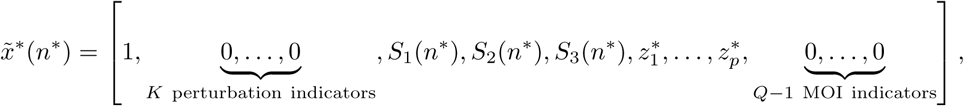

where *z^*^_j_* is the median value of technical covariate *j* across the analyzed cells. Setting perturbation-indicator columns to zero removes perturbation-specific offsets from the displayed guide-burden trend, and setting MOI indicator columns to zero evaluates the trend at the reference MOI level. With perturbation and MOI indicator columns fixed to zero, the corresponding fitted value for gene *g* is:

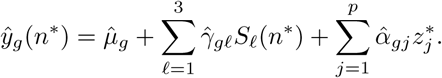

Since *z*^*^_1_, . . ., *z*^*^_*p*_ are held fixed, differences in *y*^*_g_*(*n*^∗^) across the grid are driven by the fitted spline coefficients *γ*^*_g_*_1_, *γ*^*_g_*_2_, *γ*^*_g_*_3_.

To identify genes significantly affected by the total number of detected guides, we performed a limma moderated F-test on the three spline coefficients. Specifically, for each gene *g*, we tested the null hypothesis

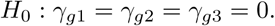

This test asks whether the fitted guide-burden curve is flat after conditioning on perturbation indicators, technical covariates, and MOI-level batch effects. P-values were adjusted for multiple testing using the Benjamini-Hochberg procedure. We found 606 genes with BH-adjusted *p* ≤ 10^−3^, which we considered burden-responsive and selected for downstream clustering and gene-signature analysis.

For these significant genes, we calculated predicted log-expression values *y*^*_g_*(*n*^∗^) across the guide-count grid. To visualize relative changes and common guide-burden trends across genes, each gene’s predicted profile was centered and scaled across the grid before heatmap visualization. This normalization removes gene-specific baseline expression and emphasizes the direction and shape of the guide-burden response.

### 1.9 Multivariate extension of reluctant interaction modeling

When considering interaction effects, we adopted a principle that one should prefer a main effect over an interaction effect if all else is equal. Yu et al. [24] introduced the sprinter (sparse reluctant interaction) algorithm that adopts this principle for univariate responses. Since Perturb-seq reads out the transcriptome with thousands of genes, we extended this approach to multivariate responses.

In the univariate case, the algorithm for reluctant interaction modeling, implemented in the R package sprintr, uses a screening criterion *I*^^^*_η_* to score an interaction:

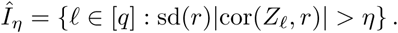

*r* = *y* − *y*^ ∈ R*^n^*^×1^ is the residual vector of the fit, where *y* is the response and *y*^ = *Xθ*^^^ is the prediction using only main effects (in Perturb-seq main effects are perturbation responses and technical covariates). cor(*Z_ℓ_, r*) is the sample correlation between the *ℓ*-th interaction variable *Z_ℓ_* and the residual *r*. *Z_ℓ_* is the *ℓ*-th column of the interaction design matrix *Z* ∈ R*^n^*^×^*^q^*, derived from the pairwise interaction between two main effects. In practice *Z, r* are centered and scaled to give *Z̃*, *r̃*. Intuitively, sd(*r*) gives the magnitude of the error we wish to explain while |cor(*Z_ℓ_, r*)| measures how well the proposed interaction is aligned with the residual.

We extended the screening criterion from a residual vector *r* ∈ R*^n^*^×1^ to a residual matrix **R** = [**r**_1_, *. . .,* **r***_k_*] ∈ R*^n^*^×^*^k^* for *k* response variables. Let **r***_j_* denote the *j*-th column of **R**, and let **R** denote the matrix obtained by centering and scaling each column of **R**. We use sd(**r***_j_*) to denote the sample standard deviation of the unscaled *j*-th residual response.

The cross-correlation matrix between all interaction terms and all residual responses, denoted by ***ρ*** ∈ R*^q^*^×*k*^, is computed via the matrix multiplication:

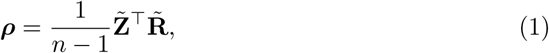

where a column of the cross-correlation matrix corresponds to the sample correlation in the univariate case *ρ_ℓ,_*_1_ = cor(*Z_ℓ_, r*_1_).

The final screening matrix **S** ∈ R*^q^*^×*k*^ is calculated element-wise as:

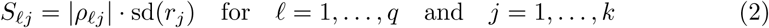

where *S_ℓj_* represents the sprinter screening statistic for the *ℓ*-th interaction term predicting the *j*-th response variable.

To apply multivariate sprinter, we first fit a main-effects Limma-voom model across 88 287 cells (*N*) and 3022 highly expressed genes (*k*). The resulting residual matrix **R** ∈ R^88287×3022^ served as the input for interaction screening. Note in Figure 1B, there are 93 305 cells because that includes cells with zero guides.

To generate the candidate interaction matrix **Z**, we constructed a primary feature matrix consisting of *p* = 55 variables, including four technical covariates (log total UMI counts, nuclear fraction, mitochondrial and ribosomal transcript fractions). The other 50 variables are indicator variables for the presence of a particular perturbation in a cell, including 45 promoter perturbations and 5 non-targeting control (NTC) guides.

Applying the multivariate sprinter score produced a 1485 × 3022 screening matrix **S**, representing the interaction strength of every pairwise feature combination for every gene 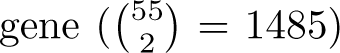. To evaluate the global impact of each interaction, we computed the mean sprinter score across all 3022 genes for each of the 1485 interaction terms.

### 1.10 Differential expression analysis with propensity score matching and marginal models

#### 1.10.1 Propensity score matching with PerturbMatch

The marginal modeling approach estimates LFCs and their standard errors after perturbation by fitting one model for each genetic perturbation. For a given perturbation *p*, we defined treatment cells as any cell containing at least one guide targeting perturbation *p*. The universe of control cells for perturbation *p* consisted of cells that contained at least one non-targeting (NT) guide and lacked any guide targeting perturbation *p*. We also excluded guides targeting the same gene at an alternative promoter from the control group. Naively comparing these treatment cells with the entire universe of control cells would create confounding from imbalanced distribution of guides per cell. Specifically, the high abundance of NT guides relative to any specific perturbation guide would cause the full control group to have fewer guides per cell compared to the treatment group. We therefore developed PerturbMatch, which stratifies by guide multiplet class (i.e. *n* = 1, *n* = 2, *n* = 3, *n* = 4, and *n* ≥ 5) to guarantee balanced distribution of guides per cell between control and treatment groups. Within each guide multiplet class, PerturbMatch applies propensity score matching [41] to further balance the distribution of technical covariates between control and treatment.

Specifically, within each guide multiplet class, we applied a two-stage procedure: (1) a weighted-GLM propensity pre-filter followed by (2) nearest-neighbor matching.

In the first stage, a logistic regression was fit to predict treatment status (i.e. whether the cell contains the guide for perturbation p) from the following technical covariates: log-transformed library size, fraction of UMIs mapping to mitochondrial genes, fraction mapping to ribosomal genes, and fraction of reads mapping to intronic regions (nuclear fraction). For cells in the *n* ≥ 5 class, guide count was additionally included as a covariate because it varies within that class. Because control cells typically outnumber treatment cells, fitting an unweighted logistic regression yields propensity scores that are shifted toward zero, making a fixed propensity score filter unreliable. To address this, we fit the logistic regression with sampling weights that equalize the effective class sizes: treatment cells received weight 1 and control cells received weight *n*_target_*/n*_NTC_ (R package `MatchIt, method=NULL, distance=“glm”`, s.weights). The resulting propensity scores are centered near 0.5, enabling a fixed filter window (e.g., [0.2, 0.8]) to reliably remove control cells whose technical profiles fall outside the support of the treatment distribution. If the weighted GLM failed to converge, we fell back to fitting an unweighted logistic regression on a balanced subsample of control cells and used the resulting model to score the full control pool.

In the second stage, the filtered control pool was passed to nearest-neighbor propensity score matching at a ratio of *K* = 5 control cells per treatment cell (R package MatchIt, method=“nearest”, distance=“glm”). If the filtered control pool was smaller than *K* times the number of treatment cells, any shortfall was filled by randomly sampling from the unfiltered controls. For quintuplets and higher (*n* ≥ 5), which were rare, the matching ratio was additionally capped at ⌊*n*_NTC_*/n*_target_⌋. If the available control pool in any class was smaller than *K* times the number of treatment cells, all available controls were retained without matching.

#### 1.10.2 Robust Limma-voom fitting for single cells

We used a modified voom pipeline adapted for single-cell RNA-seq to account for lowly expressed genes. Lowly expressed genes in single-cell transcriptomic data often have zero counts across all cells, which artificially reduces the variance to zero. With many zero-variance genes, the mean-variance trend fit by the standard voom pipeline overemphasizes the zero-variance genes, and underfits genes with moderate or high mean expression. We therefore used a robust loess smoother with a minimum spanning window, which prevents the trend from being dominated by these zero-variance, zero-expression genes. Precision weights for genes below the trend peak were further regularized to the maximum fitted standard deviation, which downweights zero-variance genes. After voom weight estimation, a standard Limma linear model was fitted per gene with empirical Bayes moderation. For standard singlet fits without multiplets, the linear model included the following covariates: a binary treatment indicator (target cells vs. non-targeting cells), log-transformed total UMI count, fraction of UMIs mapping to mitochondrial genes, and fraction mapping to ribosomal genes. For joint analysis with multiplets, the number of guides per cell was additionally included as a covariate. This covariate captures the mean passenger guide effect.

### 1.11 Quantifying information loss using neighborhood ranks

To quantify information loss across MOI, we used a neighborhood rank statistic adapted from the information imbalance framework of Glielmo et al. [30]. For two measurements *A* and *B* defined on the same set of perturbations, we first represented each perturbation *i* by a transcriptome-wide *z*-score vector, with entries log FC*/* se(log FC). We then computed pairwise Euclidean distances between perturbation profiles separately in each measurement.

Let *r^A^_ij_* denote the rank of perturbation *j* among all neighbors of perturbation *i* in measurement *A*, ordered by increasing Euclidean distance, excluding *i* itself. Thus, *r^A^_ij_* = 1 means that *j* is the nearest neighbor of *i* in measurement *A*. For each perturbation *i*, we identified its nearest neighbor in the reference measurement *A*,

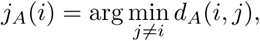

and then asked where that same perturbation *j_A_*(*i*) ranked among the neighbors of *i* in the query measurement *B*. We define the reference-to-query information loss as:

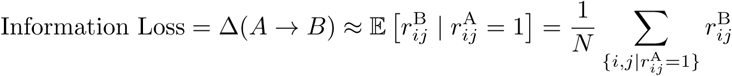

A small value indicates that each perturbation’s nearest neighbor in the reference measurement remains a close neighbor in the query measurement, whereas larger values indicate the nearest neighbor in reference is now a far neighbor in query.

Because the raw mean rank displacement lacks immediate intuition and depends on the number of perturbations in the experiment, we defined a normalized information loss *L̃_T_* for a specific test dataset *T* . This normalized loss *L̃_T_* is intuitively comparable between the 35-gene and 5000-gene Perturb-seq experiments because this loss is relative to a baseline biological and technical variation.

We calculated the fold change in loss for a test dataset *T* relative to the baseline loss observed between two measurements *R* and *R*^′^:

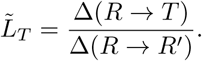

In this formulation, *L̃_T_* represents information loss as a fold change above the inherent biological and technical noise floor; a value of 1 indicates information loss equivalent to a baseline comparison. For the 35-gene targeted Perturb-seq analysis, we used low MOI as the reference (*R*) and medium MOI as the baseline comparison (*R*^′^). For the genome-wide 5000-gene analysis, we split half the singlets in the low-MOI data into a reference set and the other half in a holdout set as the baseline comparison. To reduce shared technical noise between reference and holdout, we ensured that reference and holdout cells did not come from the same 10X Genomics lanes.

### 1.12 Theoretical relationship between transcriptome-wide response and cosine similarity

Given the same genetic perturbation measured in two different experiments, we model the two high-dimensional measurements with a Gaussian additive noise model (i.e. the signal and noise components are additive). The observed standardized LFCs 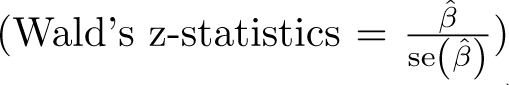 for a perturbation across *n* genes (in this paper we use roughly 3000 variable genes) can be defined as the sum of a true biological signal **s** and independent noise *ɛ* ∼ N (0, 1). For two replicates, **z**_1_ and **z**_2_, this relationship is expressed as:

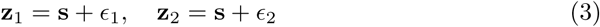

where *ɛ*_1_ and *ɛ*_2_ represent uncorrelated noise. The magnitude of the observed response is quantified by the Root Mean Square (RMS), which is related to the signal-to-noise ratio (SNR): 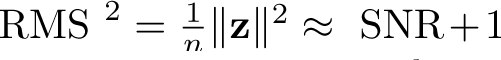. Here, the constant +1 represents the mean square (variance) of the standardized noise floor.

The theoretical expected reproducibility is defined as the expected cosine similarity *ρ* between these replicates. In high dimensions, independent noise vectors are approximately orthogonal (*ɛ*_1_ · *ɛ*_2_ ≈ 0), as is the noise relative to the signal (**s** · *ɛ* ≈ 0). Consequently, the similarity *ρ* is derived as the ratio of shared signal power (equal to SNR if standardized) to total observed power (equal to SNR + 1 if standardized):

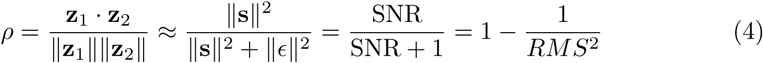

This equation relates the expected concordance between replicates to the transcriptome-wide response of a perturbation (calculated as the signal-to-noise ratio SNR = RMS^2^−1). We assessed empirical convergence to this benchmark across genetic perturbations to evaluate how different modeling scenarios reproduce the reference signal. Although the equation calculates the expected cosine similarity, it is essentially the same for Pearson correlation because the distribution of the LFC responses across the transcriptome is centered around zero, even for strong genetic perturbations. Practically, we find virtually no difference between calculating cosine similarity and Pearson correlation from LFC vectors. Consequently, the signal’s power 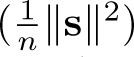 effectively matches its variance. This formulation, where the expected cosine similarity or Pearson correlation is a function of the signal-to-noise ratio, mirrors classical reliability estimation frameworks [42, 43]. In the context of modern high-dimensional genomics, our use of standardized LFCs yields a theoretical reproducibility benchmark consistent with that recently formalized by Vollenweider and Bühlmann [44] for evaluating perturbation predictions across replicates.

### 1.13 Calculating precision-recall metrics across perturbations

We calculated precision and recall by first defining a reference set, which consists of singlets in both treatments and controls. For each genetic perturbation, we identified a reference set of differentially expressed genes (DEGs) to serve as the ground truth against which the query sets were evaluated. For guide multiplet titrations, we generated query sets by gradually replacing holdout cells with a specific type of guide multiplet (e.g. doublets, triplets, or higher-order) until we reached a multiplet fraction *f* = 1, indicating pure guide multiplets of a certain class. We generated query sets on a grid of guide multiplet fractions: *f* = [0, 0.1, 0.2, 0.4, 0.6, 0.8, 1.0].

In the 35-gene targeted Perturb-seq, we used all singlets from low MOI as reference and 50% singlets across the other MOI levels as holdout. In the 5000-gene unbiased Perturb-seq, we used singlets from the Low MOI library as reference and all singlets and doublets pooled from the Medium and High MOI libraries as the holdout (the *f* = 0 baseline). In the Replogle K562 dataset, which lacks a distinct low-MOI reference library, we instead split singlets into an 80% reference and a 20% holdout set. In the Replogle dataset, rather than a continuous *f* grid, we compared three discrete scenarios: (1) the singlet holdout alone, (2) the singlet holdout combined with doublets and triplets (*n* = 4 and *n* = 6 guides per cell, corresponding to two perturbations and three perturbations per cell), and (3) multiplets alone.

For all three datasets, DEGs were defined using adaptive shrinkage (implemented in the R package ashr) [45] applied to the Limma-voom logFC estimates and their standard errors. A gene was called differentially expressed if its local false sign rate (LFSR) fell below a threshold and its posterior mean *|β|* exceeded a log_2_ fold-change threshold. For the 35-gene targeted Perturb-seq we used LFSR *<* 0.01 and *|β| >* 0.1; for the 5000-gene unbiased Perturb-seq and Replogle dataset, we used LFSR *<* 0.15 and *|β| >* 0.1. We relaxed the LFSR threshold because untargeted screens contain more perturbations with weak effects. Precision and recall were computed for each perturbation independently by comparing the DEG set of each query condition against the reference DEG set. We summarized results across perturbations by reporting the mean precision, recall, and F-score at each *f* value.

## Author Contributions

Conception: S.X. and J.Y., with input from S.M.C., W.F.F., and Jack Kamm. Experiments: J.T., L.W., and D.W., under the supervision of S.X. Data preprocessing: J.T., with help from L.W. Formal analysis: J.Y., with input from S.X., W.F.F., and Jorge Kageyama. Theory: J.Y., with input from B.B.C., O.M., W.F.F., and Jack Kamm. Writing: S.X. and J.Y.

## Acknowledgements

We thank Stefan Peidli, Linh Trinh, Brian Orcutt-Jahns, and Aviv Regev for discussions about guide-pooling strategies.

## Declarations

All authors are employees of Genentech/Roche and have equity in Roche. The funding for this project was provided by Genentech/Roche.

## Data and Code Availability

Raw and processed Perturb-seq data generated for this study have been deposited in the Gene Expression Omnibus (GEO) under accession number GSE337988. Data derived from publicly available Perturb-seq datasets are available from Zenodo at doi:10.5281/zenodo.21232999. The standalone software for propensity score matching in differential expression analysis, perturbmatch, is available on GitHub at https://github.com/Genentech/perturbmatch. Code to reproduce the analysis and figures is available on GitHub at https://github.com/Genentech/perturbmatchanalysis.

## References

[1] Bock, C., et al. High-content CRISPR screening. Nature Reviews Methods Primers 2 (2022). URL https://www.nature.com/articles/s43586-021-00093-4.

[2] Dixit, A. et al. Perturb-Seq: Dissecting molecular circuits with scalable single-cell RNA profiling of pooled genetic screens. Cell 167, 1853–1866.e17 (2016). URL https://www.cell.com/action/showFullText?pii=S0092867416316105.

[3] Adamson, B. et al. A multiplexed single-cell CRISPR screening platform enables systematic dissection of the unfolded protein response. Cell 167, 1867–1882.e21 (2016). URL https://www.cell.com/action/showFullText?pii=S0092867416316609.

[4] Jaitin, D. A. et al. Dissecting immune circuits by linking CRISPR-pooled screens with single-cell RNA-seq. Cell 167, 1883–1896.e15 (2016). URL https://www.cell.com/action/showFullText?pii=S0092867416316117.

[5] Datlinger, P. et al. Pooled CRISPR screening with single-cell transcriptome read-out. Nature Methods 14, 297–301 (2017). URL https://www.nature.com/articles/nmeth.4177.

[6] Xie, S., Duan, J., Li, B., Zhou, P. & Hon, G. C. Multiplexed engineering and analysis of combinatorial enhancer activity in single cells. Molecular Cell 66, 285–299.e5 (2017). URL https://www.cell.com/action/showFullText?pii=S1097276517301740.

[7] Replogle, J. M. et al. Mapping information-rich genotype-phenotype landscapes with genome-scale Perturb-seq. Cell 185, 2559–2575.e28 (2022). URL https://www.sciencedirect.com/science/article/pii/S0092867422005979.

[8] Huang, A. C. et al. X-Atlas/Orion: Genome-wide Perturb-seq datasets via a scalable fix-cryopreserve platform for training dose-dependent biological foundation models. bioRxiv 2025.06.11.659105 (2025). URL https://www.biorxiv.org/content/10.1101/2025.06.11.659105v1.

[9] Zhu, R. et al. Genome-scale Perturb-seq in primary human CD4+ T cells maps context-specific regulators of T cell programs and human immune traits. bioRxiv 2025.12.23.696273 (2025). URL https://www.biorxiv.org/content/10.64898/2025.12.23.696273v1.

[10] Lo, Y. H., et al. Large-scale CRISPR screening in primary human 3D gastric organoids enables comprehensive dissection of gene-drug interactions. Nature Communications 16 (2025). URL https://www.nature.com/articles/s41467-025-62818-3.

[11] Shi, T. et al. Genome-scale functional mapping of the mammalian whole brain with in vivo Perturb-seq. bioRxiv 2026.03.16.711480 (2026). URL https://www.biorxiv.org/content/10.64898/2026.03.16.711480v1.

[12] Gasperini, M. et al. A genome-wide framework for mapping gene regulation via cellular genetic screens. Cell 176, 377–390.e19 (2019). URL https://www.cell.com/action/showFullText?pii=S009286741831554X.

[13] Xie, S., Armendariz, D., Zhou, P., Duan, J. & Hon, G. C. Global analysis of enhancer targets reveals convergent enhancer-driven regulatory modules. Cell Reports 29, 2570–2578.e5 (2019). URL https://www.cell.com/action/showFullText?pii=S2211124719313956.

[14] Yao, D. et al. Scalable genetic screening for regulatory circuits using compressed Perturb-seq. Nature Biotechnology 42, 1282–1295 (2024). URL https://www.nature.com/articles/s41587-023-01964-9.

[15] Kamm, J., Yeung, J. & Forrest, B. Fishash: A contingency table approach to Perturb-seq guide assignment. bioRxiv 2026.01.22.701179 (2026). URL https://www.biorxiv.org/content/10.64898/2026.01.22.701179v1.

[16] Law, C. W., Chen, Y., Shi, W. & Smyth, G. K. voom: precision weights unlock linear model analysis tools for RNA-seq read counts. Genome Biology 15 (2014). URL https://link.springer.com/article/10.1186/gb-2014-15-2-r29.

[17] Dong, W. & Kantor, B. Lentiviral vectors for delivery of gene-editing systems based on CRISPR/Cas: Current state and perspectives. Viruses 13, 1288 (2021). URL https://www.mdpi.com/1999-4915/13/7/1288.

[18] Kim, K. H. & Lee, M. S. GDF15 as a central mediator for integrated stress response and a promising therapeutic molecule for metabolic disorders and NASH. Biochimica et Biophysica Acta 1865, 129834 (2021). URL https://www.sciencedirect.com/science/article/pii/S0304416520303457.

[19] Saxena, G., Chen, J. & Shalev, A. Intracellular Shuttling and Mitochondrial Function of Thioredoxin-interacting Protein. Journal of Biological Chemistry 285, 3997–4005 (2010). URL https://www.sciencedirect.com/science/article/pii/S0021925820810710.

[20] Liu, Y. & Bodmer, W. F. Analysis of p53 mutations and their expression in 56 colorectal cancer cell lines. Proceedings of the National Academy of Sciences 103, 976–981 (2006). URL https://www.pnas.org/doi/10.1073/pnas.0510146103.

[21] Ha, T. et al. Antisense transcription from lentiviral gene targeting linked to an integrated stress response in colorectal cancer cells. Molecular Therapy Nucleic Acids 28, 877–891 (2022). URL https://www.cell.com/action/showFullText?pii=S2162253122001421.

[22] Hrecka, K. et al. Lentiviral Vpr usurps Cul4-DDB1[VprBP] E3 ubiquitin ligase to modulate cell cycle. Proceedings of the National Academy of Sciences of the United States of America 104, 11778–11783 (2007). URL https://www.pnas.org/doi/abs/10.1073/pnas.0702102104.

[23] Zhang, S. et al. G2 cell cycle arrest and Cyclophilin A in lentiviral gene transfer. Molecular Therapy 14, 546–554 (2006). URL https://www.cell.com/action/showFullText?pii=S1525001606002395.

[24] Yu, G., Bien, J. & Tibshirani, R. Reluctant interaction modeling. *arXiv preprint arXiv:1907.08414* (2019). URL https://arxiv.org/pdf/1907.08414.

[25] Barry, T., Wang, X., Morris, J. A., Roeder, K. & Katsevich, E. SCEPTRE improves calibration and sensitivity in single-cell CRISPR screen analysis. Genome Biology 22 (2021). URL https://link.springer.com/article/10.1186/s13059-021-02545-2.

[26] Barry, T., Mason, K., Roeder, K. & Katsevich, E. Robust differential expression testing for single-cell CRISPR screens at low multiplicity of infection. Genome Biology 25 (2024). URL https://link.springer.com/article/10.1186/s13059-024-03254-2.

[27] Ahlmann-Eltze, C. & Huber, W. glmGamPoi: fitting Gamma-Poisson generalized linear models on single cell count data. Bioinformatics 36, 5701–5702 (2021). URL 10.1093/bioinformatics/btaa1009.

[28] Guo, X. et al. Recent advances in differential expression analysis for single-cell RNA-seq and spatially resolved transcriptomic studies. Briefings in Functional Genomics 23, 95–109 (2024). URL 10.1093/bfgp/elad011.

[29] Zou, H. & Hastie, T. Regularization and Variable Selection Via the Elastic Net. Journal of the Royal Statistical Society Series B: Statistical Methodology 67, 301–320 (2005). URL 10.1111/j.1467-9868.2005.00503.x.

[30] Glielmo, A., Zeni, C., Cheng, B., Csányi, G. & Laio, A. Ranking the information content of distance measures. PNAS Nexus 1 (2022). URL 10.1093/pnasnexus/pgac039.

[31] Bunne, C. et al. How to build the virtual cell with artificial intelligence: Priorities and opportunities. Cell 187, 7045–7063 (2024). URL https://www.cell.com/cell/fulltext/S0092-8674(24)01332-1.

[32] Niu, Z., et al. PerturbPlan: An analytical framework for designing Perturb-seq experiments. *bioRxiv* (2026). URL https://www.biorxiv.org/content/10.64898/2026.05.22.727199v1.

[33] Metzner, E., Southard, K. M. & Norman, T. M. Multiome Perturb-seq unlocks scalable discovery of integrated perturbation effects on the transcriptome and epigenome. Cell Systems 16, 101161 (2025). URL https://www.cell.com/cell-systems/fulltext/S2405-4712(24)00366-1.

[34] Kudo, T., et al. Scalable multimodal mapping of macrophage regulatory architecture by integrating optical and transcriptomic pooled screens. *bioRxiv* (2026). URL https://www.biorxiv.org/content/10.64898/2026.05.27.728345v1.

[35] Srinivasan, R. et al. Chemically-inducible CRISPR/Cas9 circuits for ultra-high dynamic range gene perturbation. Nature Communications 16, 1234 (2025). URL 10.1038/s41467-025-67201-w.

[36] McGinnis, C. S. et al. MULTI-seq: sample multiplexing for single-cell RNA sequencing using lipid-tagged indices. Nature Methods 16, 619–626 (2019). URL 10.1038/s41592-019-0433-8.

[37] Li, B. et al. Cumulus provides cloud-based data analysis for large-scale single-cell and single-nucleus RNA-seq. Nature Methods 17, 793–798 (2020). URL https://www.nature.com/articles/s41592-020-0905-x.

[38] Townes, F. W., Hicks, S. C., Aryee, M. J. & Irizarry, R. A. Feature selection and dimension reduction for single-cell RNA-Seq based on a multinomial model. Genome Biology 20 (2019). URL https://link.springer.com/article/10.1186/s13059-019-1861-6.

[39] Muskovic, W. & Powell, J. E. DropletQC: improved identification of empty droplets and damaged cells in single-cell RNA-seq data. Genome Biology 22 (2021). URL https://link.springer.com/article/10.1186/s13059-021-02547-0.

[40] McCarthy, D. J., Campbell, K. R., Lun, A. T. & Wills, Q. F. Scater: pre-processing, quality control, normalization and visualization of single-cell RNA-seq data in R. Bioinformatics 33, 1179–1186 (2017). URL 10.1093/bioinformatics/btw777.

[41] Rosenbaum, P. R. & Rubin, D. B. The central role of the propensity score in observational studies for causal effects. Biometrika 70, 41–55 (1983). URL 10.1093/biomet/70.1.41.

[42] Spearman, C. The proof and measurement of association between two things. By C. Spearman, 1904. The American journal of psychology 15, 72–101 (1904).

[43] Lord, F. M. & Novick, M. R. Statistical theories of mental test scores. Addison-Wesley 55–81 (1968).

[44] Vollenweider, M. & Bühlmann, P. Signal, bounds, and baselines: Principles for evaluating virtual cell perturbation models. bioRxiv 2026.04.20.719650 (2026). URL https://www.biorxiv.org/content/10.64898/2026.04.20.719650v2.

[45] Stephens, M. False discovery rates: a new deal. Biostatistics 18, 275–294 (2017). URL 10.1093/biostatistics/kxw041.

